# Structural basis for the allosteric regulation of Human Topoisomerase 2α

**DOI:** 10.1101/2020.08.23.263558

**Authors:** Arnaud Vanden Broeck, Christophe Lotz, Robert Drillien, Claire Bedez, Valérie Lamour

**Author notes:** Correspondence to Valérie Lamour.

## Abstract

The human type IIA topoisomerases (Top2) are essential enzymes that regulate DNA topology and chromosome organization. The Top2α isoform is a prime target for antineoplastic compounds used in cancer therapy that form ternary cleavage complexes with the DNA. Despite extensive studies, structural information on this large dimeric assembly is limited to the catalytic domains, hindering the exploration of allosteric mechanism governing the enzyme activities and the contribution of its non-conserved C-terminal domain (CTD). Herein we present cryo-EM structures of the entire human Top2α nucleoprotein complex in different conformations solved at subnanometer resolutions. Our data unveils the molecular determinants that fine tune the allosteric connections between the ATPase domain and the DNA binding/cleavage domain. Strikingly, the reconstruction of the DNA-binding/cleavage domain uncovers a linker leading to the CTD, which plays a critical role in modulating the enzyme’s activities and opens perspective for the analysis of post-translational modifications.

## Introduction

Type II DNA topoisomerases (Top2) are evolutionary conserved enzymes whose primordial activity is to regulate the homeostasis of DNA topology in eukaryotes and bacteria ^1^. The Top2 are involved in essential cellular processes such as DNA replication, DNA transcription, and chromosome segregation ^2^. The human Top2α isoform is highly expressed during mitosis, essential for cell division ^3^ and a biomarker for cell proliferation ^4^. As such, Top2α is a major target for antineoplastic drugs that hamper its catalytic activities ^5^.

This large homodimeric enzyme introduces a double strand break in a first DNA duplex, called G-segment, and directs the transport of a second DNA duplex, called T-segment, through the transient break in order to change the topology of a DNA crossover. The passage of the T-segment requires ATP hydrolysis and is thought to occur along with the opening and closing of several dimeric interfaces constituting molecular gates ^6,7^. The crystal structures of the ATPase and DNA binding/cleavage domains of eukaryotic Top2 have been determined and present cavities compatible with the binding of a DNA double helix ^8–13^. Biochemical and structural studies have provided evidence that the ATPase domain or N-gate, and the DNA binding/cleavage domain forming the DNA- and C-gates, are allosterically connected, a key feature of its activity ^14,15^. However, hinge regions connecting the catalytic sites of the human enzyme remain largely unexplored, hindering efforts to apprehend the quaternary organization of this enzyme and the landscape of conformations it adopts during the catalytic cycle.

In addition, the Top2 catalytic domains are flanked by C-terminal extensions that vary from one species to another ^16,17^. These domains contain nuclear localization signals and are submitted to extensive post-translational modifications that condition the cellular localization of Top2, its interactions with cellular partners and progression of the cell cycle ^18,19^. Several studies have suggested that different regions of the Top2α CTD contribute to the enzyme’s catalytic activities through DNA binding ^20–23^. In contrast with prokaryotic enzymes that harbor a pinwheel-structured CTD ^24–26^, the same region in eukaryotic enzymes presents no homology to any known fold, hence limiting structure-function analysis. It has become clear that the analysis of the molecular determinants of the enzyme’s allostery and the modulation of its activity by the CTD now requires the availability of a complete molecular structure of the Top2α.

In this work, we determined the cryo-EM structure of the full length human Top2α isoform bound to DNA in different conformations trapped by the anti-cancer drug etoposide. The structures reveal the connections between the ATPase and DNA binding/cleavage domains, allowing the identification of conserved sequence patterns in humans that control the allosteric signaling between the catalytic sites. In addition, we were able to localize the linker between the DNA binding/cleavage domain and the CTD inserting below the G-segment. We show that this region directly stimulates the Top2α catalytic activity suggesting that the bulk of the CTD domain may counterbalance this effect, potentially through post-translational modifications.

## Results

### Cryo-EM reconstructions of the hTop2α nucleoprotein complex

The full-length hTop2α was overexpressed in mammalian cells and purified using tandem affinity chromatography followed by a heparin step (see Methods and Supplementary Fig. 1). Samples were tested for enzymatic activity and mixed with asymmetric DNA oligonucleotides mimicking a doubly-nicked 30bp DNA ^11^ (Supplementary Fig. 1b and Table 2). The antineoplastic drug etoposide and AMP-PNP, the non-hydrolysable homolog of ATP, were added to the nucleoprotein complex in order to minimize the conformational heterogeneity of the sample.

The DNA-bound hTop2α complex was analyzed by single-particle cryo-EM. Six different datasets were recorded on a Titan Krios microscope yielding a total of 13,484 micrographs. Using automated picking procedure in RELION2 ^27^, a total of 1,908,092 particles were extracted from the micrographs. The dataset was curated using a combination of 2D and 3D ab-initio classifications in RELION2 and cryoSPARC ^28^, respectively, yielding a final stack of 162,332 particles (Supplementary Fig. 2). Using a low-pass filtered 3D ab-initio model generated in cryoSPARC, a first 3D reconstruction of the DNA-bound hTop2α was obtained at 6.6 Å resolution (Supplementary Fig. 3). The flexibility of the complex is visible from the 2D class averages showing the dimerized ATPase domain adopting different positions relative to the DNA-binding/cleavage domain (Supplementary Fig. 2b,d).

To isolate the different conformations and improve the overall resolution, we used a combination of global and local approaches to deconvolute the structures of the hTop2α complex. Using 3D focused refinements, reconstructions of the DNA-binding/cleavage domain were generated for two different states. State 1 was solved at 4.2 Å using 57,976 particles and State 2 was solved at 5.0 Å using 34,922 particles (Supplementary Fig. 3). Each particle stack was submitted to 3D-refinement without mask to yield two reconstructions of the entire complex in State 1 at 5.6 Å resolution, and in State 2 at 7.2 Å resolution (Supplementary Fig. 3). Due to the flexibility of the ATPase domain, the densities of the linkers with the DNA binding/cleavage domain were not well resolved in the cryo-EM maps. To get information on this region, a 3D-focused classification was performed on the ATPase domain yielding one class comprising 36,610 particles with a well-resolved connection between the two functional domains which was refined at 7.6 Å resolution (Supplementary Fig. 3).

### Model building of the hTop2α complex in different states

Near-atomic resolution features of the 4.2 Å map in State 1 allowed us to fit, build and refine the atomic model of the hTop2α DNA-binding/cleavage domain in complex with DNA and etoposide ^29^ (Figure 1a-c, Supplementary Fig. 5). Strikingly, we were also able to identify EM density for the etoposide molecule, intercalating in the DNA duplex between positions −1 and +1 (Figure 1d), with similar protein and DNA contacts as previously reported in crystallographic structures (Supplementary Fig. 7) ^10,29^.

**Figure 1.**
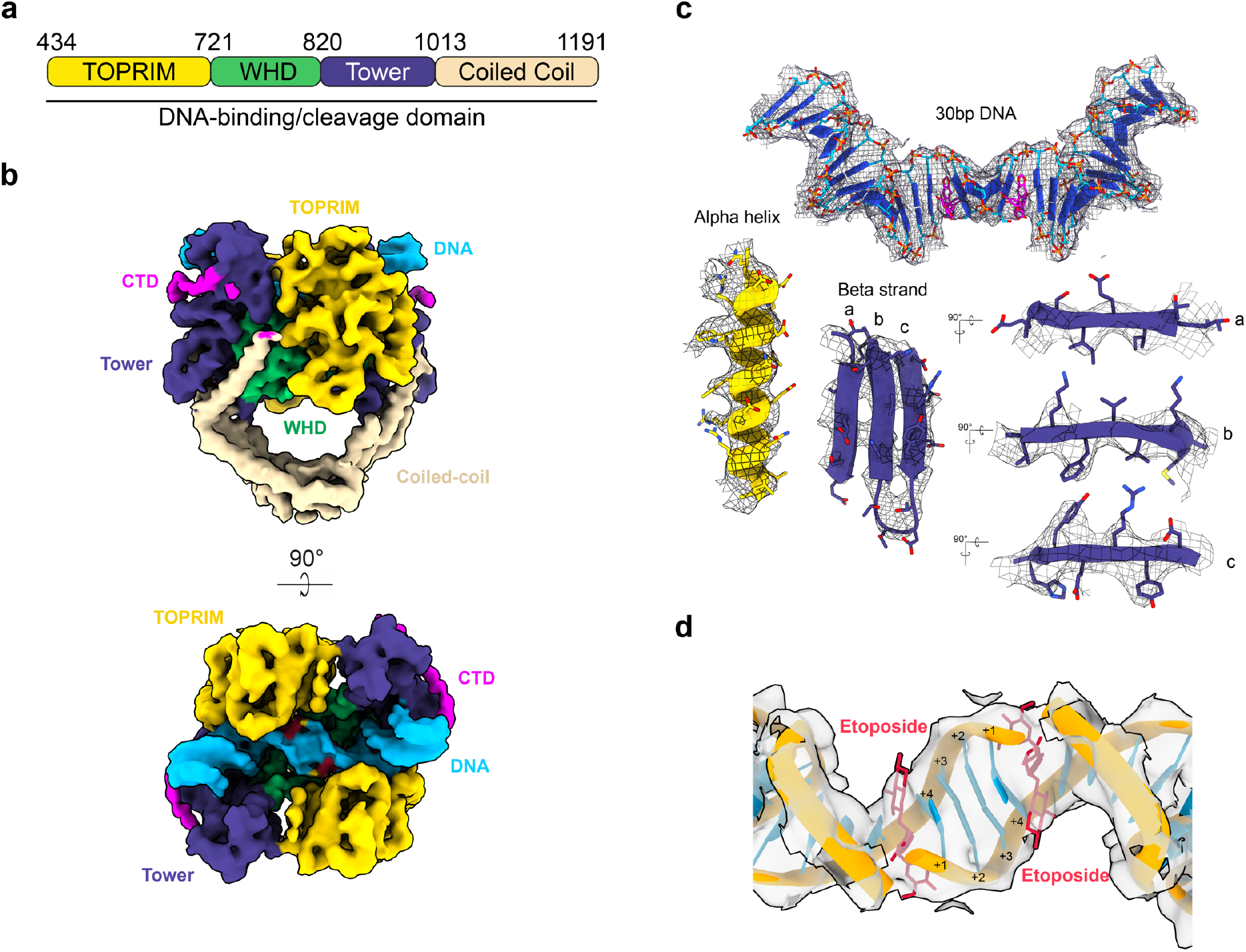
Cryo-EM structure of the etoposide-poisoned DNA-binding/cleavage domain. **a**. Schematic representation of the hTop2α DNA-binding/cleavage domain. **b.** Cryo-EM structure of the DNA-binding/cleavage domain in State 1 solved at 4.2Å resolution. The structure is colored as in a, the DNA is colored in blue and a visible fragment of the CTD is colored in magenta. **c.** Near-atomic resolution features of the 4.2Å cryo-EM map. The 30bp DNA and the corresponding EM density are shown in the upper part. An alpha helix with well-resolved side chain densities of aromatic and basic residues is shown on the left. Beta-strands with well-resolved side chain densities of aromatic and basic residues are shown on the right. **d.** EM density around etoposide and the intercalated DNA bases is shown. The +1 position is not phosphorylated to trap the DNA-bound hsTop2a in a cleavage complex-like state.

The resulting DNA-binding/cleavage domain model together with the crystal structure of the ATPase domain bound to AMP-PNP ^8^ were then combined to refine the complete atomic structure of the full-length hTop2α in State 1 (Figure 2). The 27-aa linkers between the two domains, missing from the crystal structure of the isolated ATPase domain, were built as alpha helices based on secondary structure predictions and using well-defined linker densities of the cryo-EM map obtained by focused classification on the ATPase domain (Supplementary Fig. 3 and 6). Finally, the atomic model of the DNA-binding/cleavage domain in State 2 and the corresponding full-length hTop2α in State 2 were refined using State 1 model as a reference.

**Figure 2.**
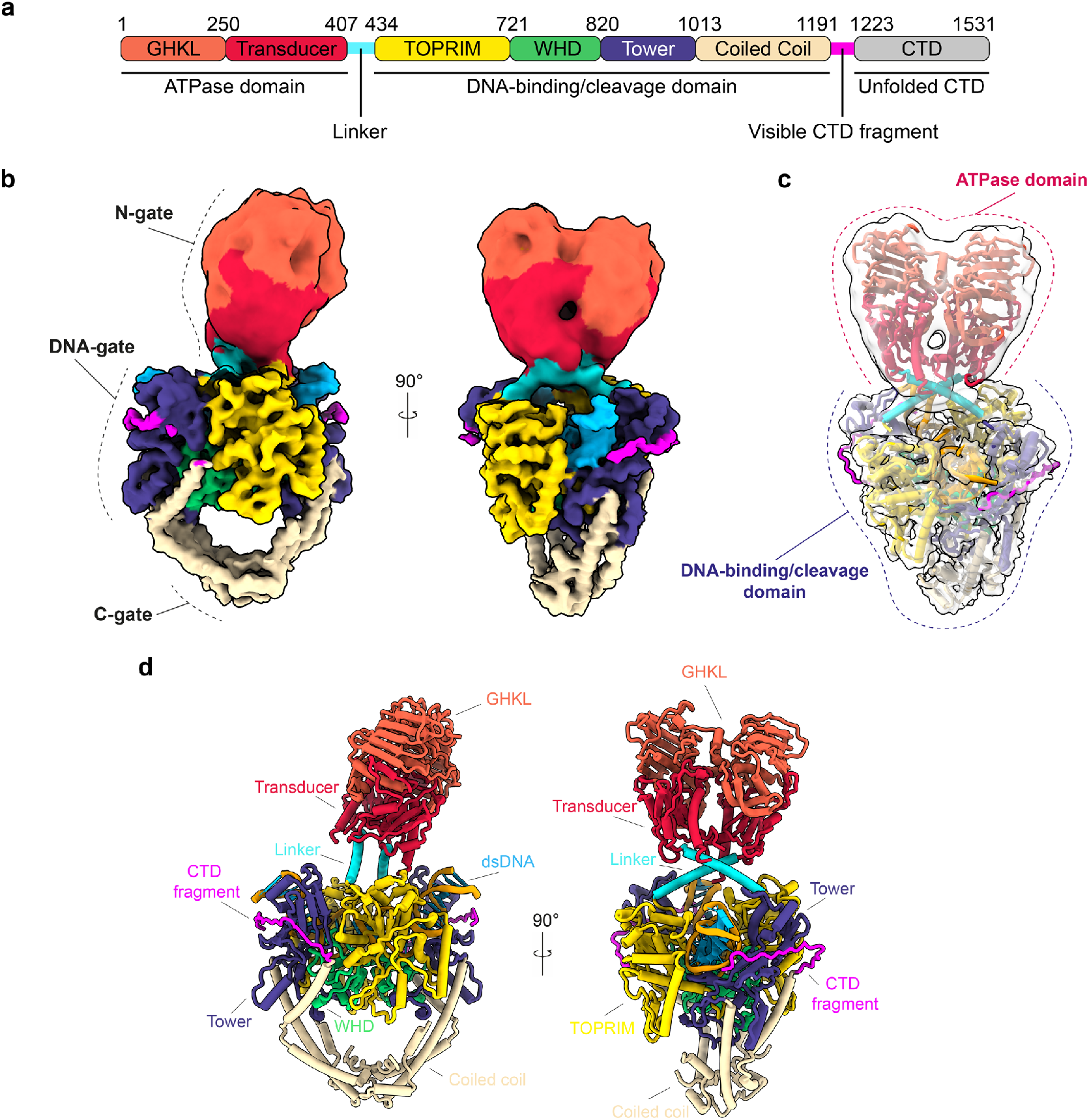
Molecular model of the DNA-bound Human Top2α. **a.** Schematic representation of the hTop2α domains. Human Top2α is composed of two identical subunits assembling in an active homodimeric form. **b**. Composite cryo-EM map of the full Human Top2α with DNA-gate in State 1. **c.** Atomic model fitted inside the cryo-EM map. Each domain is coloured as in a. **d.** Atomic model of the full human Top2α with the same colour coding.

### Analysis of the hTop2α conformations

The complete architecture of the hTop2α reveals the intertwined arrangement of the two subunits, a feature that was deduced from the yeast enzyme crystal structure and recently observed in the cryo-EM models of the prokaryotic Topo II ^12,30^ (Figure 2d and Figure 3d-f). The dimeric ATPase domain sits in a ~95°orthogonal orientation above the DNA-gate, similar to what was previously observed with the yeast Top2 (~90° orientation) ^12^ (Figure 3d-f). The structure is asymmetric with the ATPase domain slightly bent (∼5°) in both orthogonal plans (Figure 3d-e). Compared to the yeast Top2 structure, the ATPase domain is positioned 15 Å above the DNA-gate. Interestingly this creates a cavity large enough to accommodate a T-segment on top of a G-segment (Figure 3d-e).

**Figure 3.**
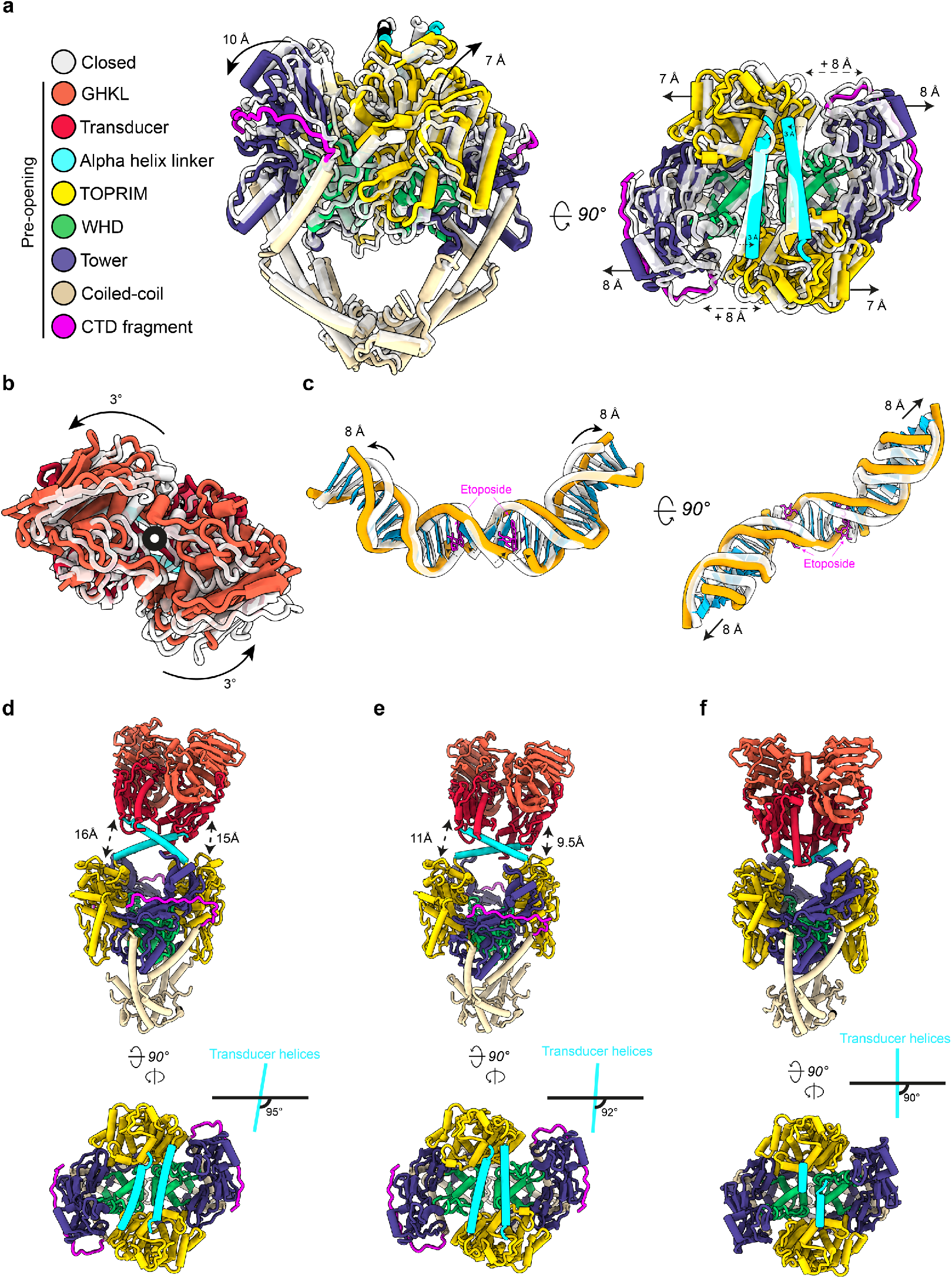
Structural changes associated with G-segment opening and allosteric connections between the N-and DNA-gates. **a**. Superimposition of DNA-binding/cleavage domain structures in State 1 (closed) (grey) and State 2 (pre-opening) (colored based on the legend). TOPRIM domain and Tower domain are moving away from 8 Å in the pre-opening state. **b**. A 3° rotating movement of the ATPase domain (opposite to the intertwining) is observed during the transition from closed to pre-opening conformation of the DNA-gate. **c**. During the transition, the DNA is unwound and stretched by 8 Å. **d**. hTop2α with DNA-gate in closed state: the ATPase domain is bent by ~5° with respect to the vertical axis and also by ~5° in the orthogonal plane, positioning the ATPase domain at a distance of ~15Å of the DNA-gate (top). The ATPase domain transducer alpha-helices in cyan form a ~95° angle with the DNA-gate (bottom). **e**. hTop2α with DNA-gate in pre-opening state: the ATPase domain is bent by ~8° with respect to the vertical axis and also by ~5° in the orthogonal plan, positioning the ATPase domain at a distance of ~10Å of the DNA-gate (top). The ATPase domain transducer alpha-helices in red form a ~92° with the DNA-gate (right). **f.** Yeast Top2 with DNA-gate in closed state: the ATPase domain is straight in both orthogonal plans, positioning the ATPase domain in close contact with the DNA-gate (top). The ATPase domain transducer alpha-helices in red form a ~90° with the DNA-gate (bottom).

In State 1, the TOPRIM and Tower domains are positioned upward and tightly bound to the G-segment which is highly bent. This state corresponds to a closed cleavage complex as observed in the structure of the DNA-bound hTop2α DNA-binding/cleavage domain crystallized without etoposide, with the exception of the C-gate being closed in our structure ^11^. In State 2, the TOPRIM domain is moved upward by 7 Å while the Tower domain is moved away from the TOPRIM domain by 8Å (Figure 3a). The physical separation of the TOPRIM and tower domains induces the stretching and unwinding of the G-segment by 8 Å in both directions (Figure 3c), positioning the DNA-binding/cleavage domain in a pre-opening conformation as observed in the crystal structure of DNA/etoposide-bound hTop2α DNA-binding/cleavage domain ^29^. In this conformation, the TOPRIM and tower domains are separated by ~20Å, which could contribute to the formation of the cavity compatible with the presence of a T-segment before transport through the G-segment.

Since the 2 structures solved in different states are bound to AMP-PNP and etoposide, the complexes are trapped in a form precluding the opening of the G-segment. However, the hTop2α DNA binding/cleavage domain is still able to oscillate between the closed and pre-opening states in presence of etoposide, despite the fact that the G-segment base pairs remain annealed (see supplemental analysis). This conformational oscillation has also been observed in different bacterial DNA gyrase complexes bound to antibiotics ^30,31^ (Figure 3, Supplementary Fig. 3 and 8).

We also found that the ATPase domain adopts different inclinations depending on the conformation of the DNA-binding/cleavage domain. When the DNA-binding/cleavage domain is a closed state, the ATPase domain is bending over by 5° relative to the dimeric symmetry axis. In the pre-opening state, the ATPase domain bends over by about 15° (Supplementary Fig. 2d and 3). These tilting movements are inherent to the flexibility of the ATPase domain which sits on top of a cavity only partially occupied by the DNA G-segment. It is most likely that the presence of a fully formed DNA crossover would constrain the tilting range of the N-gate.

The conformational changes observed in the DNA-gate are however directly correlated to rotations and translational movements of the ATPase domain (Figure 3a,b). Conformational changes of the TOPRIM domains in the DNA-gate modify the distance between residues N433 of each monomer, which are connected to the alpha helices of the ATPase transducers. In the closed state, the distance between the two residues N433 is 49.3 Å while the distance is shortened to 46.4 Å in the pre-opening state. Consequently, this translational movement of 3 Å in the orthogonal plane of the DNA gate induces the rotation of the ATPase domain by 3° counterclockwise, opposite to the intertwining (Figure 3b). It also forces the ATPase domain to come closer to the DNA-gate prefiguring a conformation that would position the T-segment in the newly formed groove between the TOPRIM and tower domains (Figure 3e). Altogether the rotation of the N-gate that correlates with the oscillation of the DNA-gate can be associated with a corkscrew mechanism that we can extrapolate from the two overall conformations of the full-length catalytic core (Supplementary Movie 1).

### Structure-based analysis of molecular determinants of allosteric transitions

Although the N-gate is not required for G-segment cleavage, the DNA gate *per se* is not able to open unless ATP binds to the N-gate ^32^. These findings support a model that implies a direct coupling between the ATP binding/hydrolysis and the DNA-gate opening through the 27-aa alpha helices linkers.

To analyze the molecular determinants of this allosteric mechanism, we performed a sequence analysis of the TOP2 protein from 37 species, including Top2α and Top2β from 5 eukaryotic organisms, from unicellular organisms to Human (Figure 4a). A conservation profile of the 27-aa linker, predicted to fold as an alpha helix, was calculated from 32 sequences out of the 37 species, including only multicellular animals, revealing two highly conserved consensus motifs; W_414_xxF_417_K_418_ and K_425_K_426_C_427_ (Figure 4b). The two aromatic residues W414 and F417 form a hydrophobic patch between the linkers, which could contribute to the stability of their interaction (Figure 4d). Lysine 418 is strictly conserved and is close to the K-loop (342-344), that was shown to be involved in DNA sensing in the yeast enzyme ^12^. Residues K425, K426 and C427 are also highly conserved and are located towards the end of the linker helices, at the entrance of the TOPRIM domain (Figure 4d).

**Figure 4.**
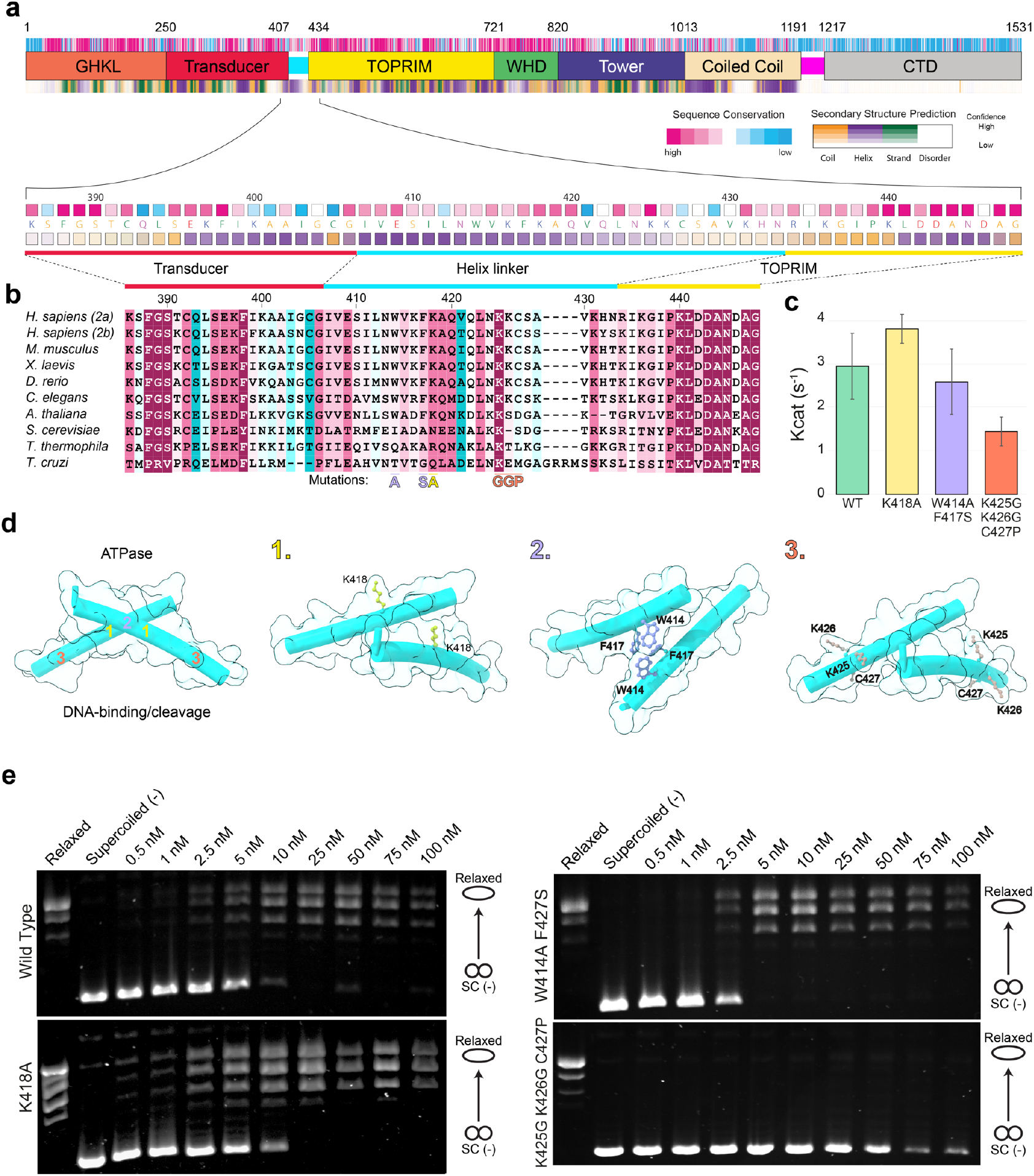
Analysis of the hTop2α allosteric regulation mediated by the linker connecting the N-gate to the DNA-gate. **a.** Domain organization, sequence conservation and secondary structure prediction for hTop2α. Below is a zoom view of the linker joining N-gate to DNA-gate showing the sequence at a residue level. **b.** Multiple sequence alignment focused on the linker region. Only 10 model organisms species of out of the 41 species used to calculate the sequence conservation are displayed. **c.** ATP hydrolysis activity of the wild-type, K418A, W414A-F417S and K425G-K426G-C427P hTop2α. Kcat was measured in triplicates at 37°C with 1mM ATP in presence of 6nM linear DNA (pUC19). **d.** Spatial localization of the mutated residues on the linker joining N-gate to DNA-gate. **e.** Relaxation activity of the wild-type, K418A, W414A-F417S and K425G-K426G-C427P hTop2α. Protein concentrations are listed in nM of holoenzyme.

To assess the contribution of these residues in the allosteric regulation of the human enzyme, we designed three hTop2α mutants: K418A to remove the positively charged side chain, W414A-F417S to remove the hydrophobic patch and K425G-K426G-C427P to disrupt the alpha helix fold. We tested their ability to perform DNA relaxation and DNA cleavage and their ATPase activity in comparison with the wild-type enzyme.

The K418A mutant displays DNA relaxation, cleavage and ATPase activities in the same range as the WT enzyme (Figure 4c,e and Supplementary Fig. 10). The W414A-F417S mutant shows a slight increase in relaxation activity but with cleavage and ATPase activities similar to the WT (Figure 4c,e, Supplementary Fig. 10). This increased DNA relaxation activity may be attributed to the disappearance of the hydrophobic patch maintaining the two linkers locked together. If this interaction is weakened, this could potentially facilitate the opening of the transducer helices during DNA-gate opening and transport of the DNA T-segment, thus slightly increasing the overall DNA relaxation activity.

On the contrary, introduction of the triple mutations K425G-K426G-C427P results in an impaired relaxation activity compared to the WT enzyme (Figure 4e). Similarly, the ATPase activity is reduced 2-fold (Figure 4c). However, the triple mutant enzyme is still able to cleave DNA, but to a lesser extent (Supplementary Fig. 10). The disruption of the helical structure may loosen the linkers, weaken their rigidity and attenuate the transmission of the signal through the linker. Overall, the different mutations show that conduction of the allosteric signal between the N-gate and the DNA-gate is finely tuned by sequence specific motifs in the linkers.

### Regulation of the catalytic activity by the C-terminal domain

Although we used the complete hTop2 sequence for our structural analysis by cryo-EM, we were not able to observe any density belonging to the bulk of the CTD in the 2D classes, nor in the 3D reconstructions. Secondary structure prediction on the CTD sequence (residues 1191-1531) suggests that this domain is disordered (Figure 5a). We confirmed this prediction by analyzing the CTD structure in solution using Small Angle X-ray Scattering (SAXS). The Kratky plot derived from the scattering curve showed a plateau for high scattering vector q-values, typical of unfolded proteins, in contrast with the Gaussian curve observed in globular domains ^33^ (Supplementary Fig. 9). However, analysis of the 4.2 Å EM map of the DNA binding/cleavage domain revealed an additional EM density that could be attributed to the beginning of the CTD domain (residues 1192-1215) (Figure 5c and Supplementary Fig. 5g). This region of the CTD begins at the end of the terminal coiled-coil alpha helix of the DNA binding/cleavage domain on residue 1192, stretches along the Tower domain and extends under the G-segment major groove pointing in an orthogonal direction from the DNA gate (Figure 5c).

**Figure 5.**
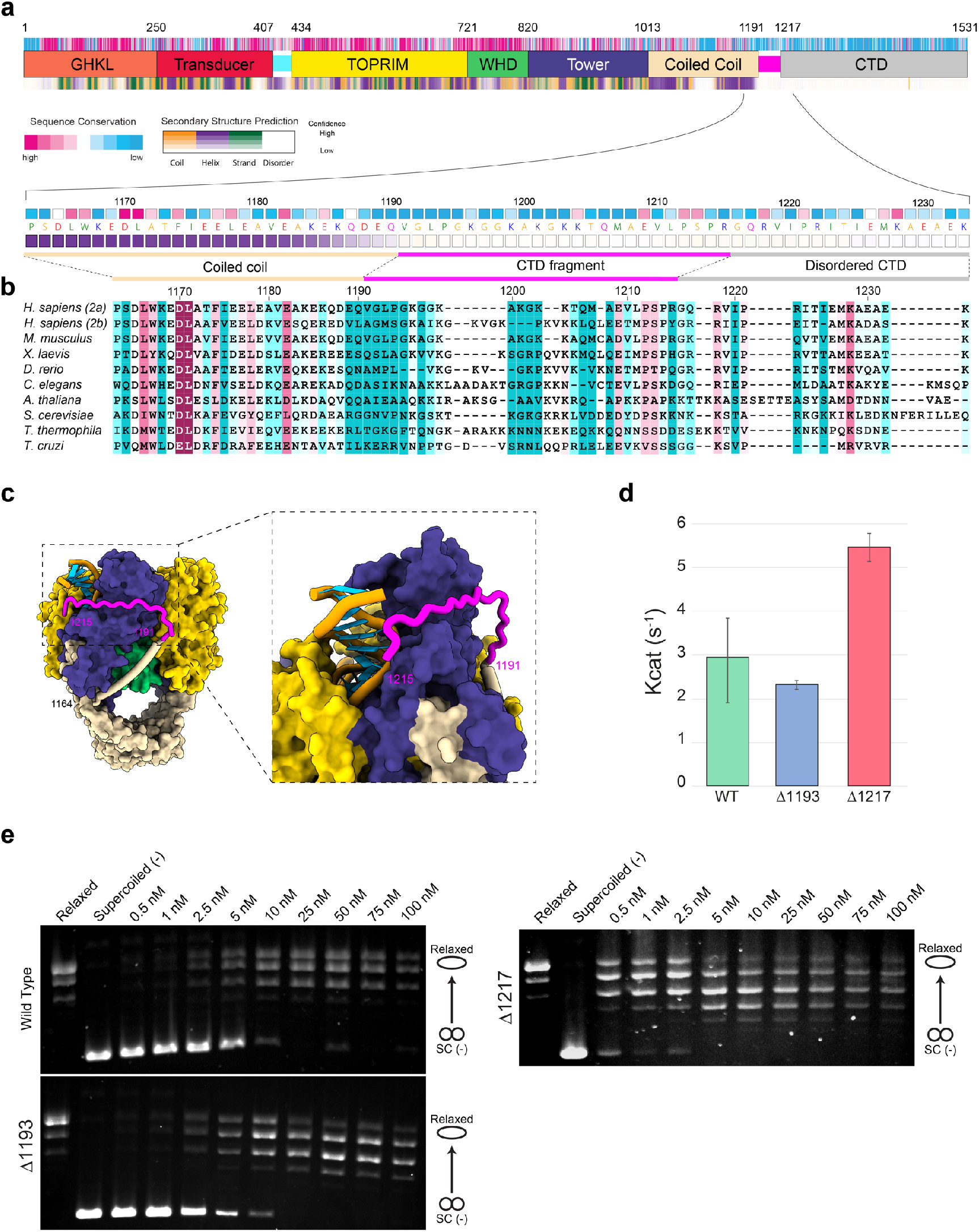
Analysis of the connection between the DNA gate and the CTD domain. **a.** Domain organization, sequence conservation and secondary structure prediction for hTop2α. Below is a zoom view of the CTD fragment visible in the EM reconstruction showing the sequence at a residue level. **b.** Multiple sequence alignment focused on the CTD fragment region. Only 10 model organisms species of out of the 41 species used to calculate the sequence conservation are displayed. **c.** Orthogonal views of the DNA-binding/cleavage domain in closed state. The CTD domain is highlighted in magenta. The inset shows a zoom on the region of the CTD which comes in close vicinity of the DNA G-segment. **d.** ATP hydrolysis activity by wild-type, Δ1193 and Δ1217 hTop2α. Kcat was measured in triplicates at 37°C with 1mM ATP in presence of 6nM linear DNA (pUC19) **e.** Comparison of relaxation activity by wild-type, Δ1193 and Δ1217 hTop2α. Protein concentrations are listed in nM of holoenzyme.

To our knowledge, the linker leading to the CTD has not been previously observed in a eukaryotic Top2 structure. The CTD has been shown to confer specific functions and DNA topology preferences to the human isoforms that differ in this region ^34,35^. It also undergoes multiple post-translational modifications that regulate its cellular distribution and activity throughout the cell cycle ^2^. Several studies have explored how the CTD could modulate the catalytic activities and DNA binding properties of the Top2 ^21,36^. However, most studies used a construct of Top2 ending before, or at position 1192, therefore not accounting for the contribution of this linker region that is in close proximity with the G-segment ^37^. To assess the contribution of this region in the catalytic activities, we designed hTop2α constructs with a complete deletion of the CTD (Δ1193) or a partial CTD truncation (Δ1217) (Figure 5). The hTop2α lacking the complete CTD (Δ1193) showed similar relaxation and ATPase activities than those of the WT enzyme (Figure 5d,e, and Supplementary Fig. 10), as also observed in previous studies ^36^. The mutant (Δ1193) is however slightly impaired in its cleavage activity compared to the WT in presence of etoposide (Supplementary Fig. 10). A similar effect of a larger Δ1175 CTD deletion was already observed with high concentrations of drug, showing that loss of the CTD decreases etoposide cleavage ^36^. On the contrary, the relaxation activity of the hTop2α with the partial CTD truncation (Δ1217) was increased ~10-fold compared to the wild-type or Δ1193 enzymes. Remarkably, its ATPase activity was also increased by ~2-fold (Figure 5d,e) and the cleavage activity of the Δ1217 was also strongly increased (Supplementary Fig. 10).

In the partial CTD truncation, the portion of the CTD ranging from residue 1192 to residue 1217 is in close vicinity of the G-segment and seems to act as a positive regulator of the strand passage process, which appears to stimulate the relaxation activity.

## Discussion

The cryo-EM structures of the entire hTop2α, reveal how the ATPase domain is spatially connected to the DNA-binding/cleavage domain conformations. The alpha helices that were missing in crystal structures of the human ATPase domain, are reminiscent of the transducer helices found in procaryotic Top2 enzymes but differ in sequence and in the relative orientation of their crossing, which narrows down the cavity of the N-gate (Figure 4 and supplementary Fig. 6). Previous crystal structures of type 2 topoisomerases isolated domains or cryo-EM reconstructions of the bacterial homolog DNA gyrase reported distinct DNA/binding domain conformation but so far without connection with the position of the ATPase domain^15,30,38^. Our structures highlight how subtle movements in the DNA-gate propagate to the N-gate through networks of interactions mediated by conserved motifs, in particular in the transducer helices. Although AMP-PNP, a non-hydrolysable analogue of ATP, was used to dimerize the ATPase domains, this shows that conformational transitions can occur within the same sample. Indeed, it was shown that AMP-PNP can support a single, complete round of DNA passage, but that the ATPase domains remain dimerized, preventing further rounds of activity ^39,40^. Structures with other nucleotides would provide further insights in the conformational range that link rotation of the N-gate to opening of the DNA gate. Although the isolated N-gate of prokaryotic enzymes have been shown to bind a DNA duplex within its cavity ^41^, it is unlikely that the human enzyme could accommodate a DNA double helix without subsequent conformational rearrangement and partial opening of the transducer helices.

In addition, the near-atomic resolution structure of the DNA-binding/cleavage domain reveals a part of the CTD domain which was not previously observed and is positioned nearby the G-segment inflection points on each side of the enzyme. The curvature of the DNA was shown to be important for the selection of cleavage sites ^42^. The particular localization of the CTD linker may structurally favor the curvature of the G-segment, stimulating DNA cleavage as shown in the cleavage assays, and favoring strand passage. We also show that this linker preceding the CTD domain enhances the catalytic activities of the hTop2α. As the Δ1192 mutant devoid of CTD shows a similar relaxation activity as the full length hTop2α, this suggests that the bulk of the CTD domain counterbalances the stimulating effect of the linker.

Interestingly the orientation of the linkers suggests the CTD domain could be positioned in an orthogonal direction relative to the plane of the Top2 catalytic domains. On the contrary, in prokaryotic enzymes, the linkers of the CTD domains are positioned in the same plane ^24,30^. The CTD linker sequence of the Top2α displays a high content in positively charged residues, as well as several conserved residues at positions 1209 and 1212-1214 also found in the Top2β isoform and other eukaryotic species (Figure 5b). This suggest that the CTD linker constitutes a common structural feature of the multicellular eukaryotic Top2, independently of the conservation of the rest of their CTD.

Interestingly, Serine 1213, located at the end of the linker, has been found to be subjected to mitotic phosphorylation and contributes to localization of the Top2α to the centromere ^43–45^. Such post-translational modification could regulate the binding of this CTD portion to the G-segment in order to modulate the relaxation activity of the hTop2α activities during the cell cycle. The Top2 activities are indeed associated with cellular complexes regulating the genome organization ^46^. The chromatin tether, a specific sequence within the hTop2α CTD, was shown to interact with histone tails in chromatin structures ^18^. The path of the CTD linker along the catalytic core of the enzyme indicates that the CTD may be positioned in a different orientation compared with the prokaryotic homologs, and may favor its binding to nucleosome structures in the eukaryotic genome.

## Acknowledgements

We thank Julio Ortiz and Gabor Papai for their help with data collection. This work was supported by the Fondation ARC, the Association Alsace contre le Cancer and the grant ANR-10-LABX-0030-INRT (managed by the Agence Nationale de la Recherche under the frame programme Investissements d’Avenir ANR-10-IDEX-0002-02). The authors acknowledge the support and the use of resources of the French Infrastructure for Integrated Structural Biology FRISBI ANR-10-INBS-05 and of Instruct-ERIC. Computational resources were provided by the Méso-centre de Calcul (University of Strasbourg).

## Contributions

A.V.B. and V.L. conceived the study and designed the experiments; A.V.B., C.L., R.D. and C.B. performed the experiments; A.V.B. performed the cryo-EM data collection, processed the cryo-EM data and built the atomic models; A.V.B., C.L. and V.L. analyzed and interpreted the data; A.V.B., C.L. and V.L. wrote the manuscript.

## Corresponding author

Correspondence to Valérie Lamour.

## Methods

### Protein expression and purification

The sequence coding for the full length human Top2α (1-1531) was inserted into a modified pVote0GW vector depleted of *attB1* sequence and containing an N-terminal Twin-strep tag and a C-terminal 10 His-tag. A selective marker and a portion of the viral hemagglutinin gene sequence (HA) present in the plasmid allowed selecting recombinant MVA-T7 viruses holding the *hTop2α* construct under the dependence of a T7 promoter. Generation and selection of the recombinant viruses was performed as described previously ^47^. Prior to overexpression, 600mL of BHK21 C13-2P cells in suspension (10^6^ cells/ml) were infected with approximately 0.1 PFU/cell of recombinant virus in cell culture medium (GMEM, 10% FCS, 1.5 g/L BTP, 50 μM Gentamycin) and incubated at 37°C. After 48h, the infected cells were mixed with 6L of uninfected cells at 10^6^ cells/ml and a 1:10 ratio (v/v), respectively. Overexpression was directly induced by the addition of 1mM IPTG followed by an incubation of 24h at 37°C. Cells were harvested and resuspended in Lysis buffer (25 mM Hepes, 500 mM NaCl, 500 mM KCl, 1 mM MgCl_2_, 20 mM imidazole, 10% v/v glycerol, 2.5 mM beta-mercaptoethanol, 0.5 mM PMSF, 0.5 mM Pefabloc, protease inhibitor cocktail (Roche), pH 8.0) and lysed with 3 cycles of high pressure disruption using an EmulsiFlex-C3 apparatus (Avestin) at 1500 bars. The full length hTop2α was first purified by a tandem affinity chromatography on a manually packed XK 26/20 column (Pharmacia) with Chelating Sepharose 6 Fast Flow resin (GE healthcare) bound to Ni^2+^ ions followed by a StrepTrap HP column (GE healthcare). Elution from the chelating resin was performed using 250mM imidazole pH 8.0 added to the Lysis buffer and allowed the protein to bind to the StrepTactin Sepharose resin. The protein was washed with 25 mM Hepes, 200 mM NaCl, 200 mM KCl, 1 mM MgCl_2_, 10% v/v glycerol, 2 mM DTT, pH 8.0 and eluted with the same buffer supplemented with 3mM Desthiobiotin (Sigma). Twin-strep and His tags were removed by the addition of P3C (Precission protease) at 1:50 ratio (w/w) and incubated overnight at 4°C. The cleaved protein was then loaded on a HiTrap Heparin HP column (GE Healthcare). Elution was performed by a single step using 25 mM Hepes, 400 mM NaCl, 400 mM KCl, 1 mM MgCl_2_, 10% v/v glycerol, 2 mM DTT, pH 8.0. After the purification process (Supplementary Fig. 1a), 20 mg of the full length hTop2α were obtained from 6L of BHK21 C13-2P cell cultures. About 10 to 15% of the protein sample shows some degradation depending on the protein batch, as previously observed ^43^. Western blot analysis using TOP2A antibody PA5-46814 (Thermo Fischer Scientific) shows that the C-terminal domain tends to be cleaved off during protein purification despite the use of protease inhibitors (Supplementary Fig. 1a). However, the majority of the sample is constituted by full-length protein forming an intact homodimer that can be stabilized to form complexes with DNA prior to cryo-EM sample freezing (Supplementary Fig. 1c).

### Nucleic acid preparation

A doubly nicked 30 bp DNA duplex was reconstituted using 2 asymmetric synthetic oligonucleotides obtained from Sigma-Aldrich. Sequences for the single strand 17bp (5’-CGCGCATCGTCATCCTC-3’) and the single strand 13bp (5’-GAGGATGACGATG-3’) were chosen as described in ^11^. Briefly, the nucleic acids were dissolved in DNAse-free water at 1 mM concentration. To assemble the double stranded DNA, each oligo was mixed at 1:1 molar ratio, annealed by incubating at 95°C for 2 min and then decreasing the temperature by 1°C every 1 min until reaching 20°C. The annealed doubly nicked DNA duplex was then buffer-exchanged in Hepes 20 mM pH 7.5 with a BioSpin 6 column (BioRad).

### Complex formation for cryo-EM

Purified hTop2α was mixed with the 30 bp dsDNA at 1:1 molar ratio to obtain a 20 μM final concentration of protein and DNA. The mixture was incubated for 10 min at 37°C. Then, etoposide (Sigma) was added to reach a final concentration of 0.5 mM. The subsequent mixture was incubated for 10 min at 37°C. Finally, ADPNP was added to the complex at a final concentration of 0.5 mM. The fully reconstituted complex was further incubated 30 min at 30°C.

### BS3 Cross-linked hTop2α-DNA-Etoposide-ADPNP complex

The purified hTop2α is unstable in buffer conditions under low salt conditions. Therefore, a chemical stabilization by BS3 was necessary for further cryo-electron microscopy analysis. To determine the optimal concentration of BS3 allowing a complete stabilization of the complex without inducing aggregates, the newly formed complex was incubated with different concentration of BS3 and analyzed by SDS-PAGE. Briefly, BS3 (Thermo Fischer Scientific) was freshly resuspended in filtered DNAse-free water at 25 mM stock concentration. Rapidly after BS3 preparation, the complex was incubated for 30 min at 30°C with 0.25 mM up to 5 mM BS3 and the crosslinking was quenched by adding Tris-HCl, pH 7.5, to 50 mM. The optimal concentration was determined at 1 mM (Supplementary Fig. 1c). The cross-linked complex was centrifuged at 20,000g for 30 min at 4°C to remove remaining aggregates.

### Buffer exchange of the complex in optimal Cryo-EM buffer

The complex was first dialyzed against 200 mL of intermediate buffer (20 mM Hepes, 200 mM KAc, 200 mM Na-Glu, 5 mM MgAc, 0.5 mM TCEP, pH 7.5) for 2h at 4°C using Slide-A-Lyzer MINI Dialysis Units (7000MWCO) (Thermo Fischer Scientific). Then, a second dialysis was performed in 200 mL of the final cryo-EM buffer (20 mM Hepes, 100 mM KAc, 50 mM Na-Glutamate, 5 mM MgAc, 0.5 mM TCEP, pH 7.5) for 4 h at 4°C. Finally, 8 mM CHAPSO (Sigma-Aldrich) was added to the dialyzed complex to prevent adsorption of the particles to the air/water interface ^48^. The sample was centrifuged 2h at 16,000 g to remove potential aggregates.

### Cryo-EM grid preparation

Quantifoil R-1.2/1.3 300 mesh copper grids were glow-charged for 20s prior to the application of 4 μl of the complex. After 30 s of incubation, the grids were plunge-frozen in liquid ethane using a Vitrobot mark IV (FEI) with 95 % chamber humidity at 10°C.

### Electron microscopy

Cryo-EM imaging was performed on a Titan Krios microscope operated at 300 kV (FEI) equipped with a Gatan K2 Summit direct electron camera (Gatan), a Gatan Quantum energy filter, and a CS corrector (FEI). Images were recorded in EFTEM nanoprobe mode with Serial EM ^49^ in super-resolution counting mode with a super resolution pixel size of 0.55 Å and a defocus range of −1 to −3.2 μm. Six datasets were collected with a dose rate of 6 to 8 e^−^/pixel/s (1.1 Å pixel size at the specimen) on the detector. Images were recorded with a total dose of 50 electrons/Å^2^, exposure time between 7 to 10 s and 0.2 to 0.25 s subframes (35 to 50 total frames).

### Data processing

Processing of each data set was done separately following the same procedure until the 3D refinements where particles were merged. The super-resolution dose-fractionated subframes were gain-corrected with IMOD ^50^ and binned twice by Fourier-cropping, drift-corrected and dose-weighted using MotionCor2 ^51^ yielding summed images with 1.1 Å pixel size. The contrast transfer function of the corrected micrographs was estimated using GCTF v1.06 ^52^. Thon rings were manually inspected for astigmatism and micrographs with measured resolutions worse than 5 Å were excluded. Particles were automatically picked by template matching in RELION2 ^27,53^. To generate the templates, around 6000 particles were manually picked on micrographs from the first data set using EMAN2 ^54^. Then, the manually picked particles were subjected to 2D classification in RELION2 and the best class averages were used as templates for subsequent automatic picking procedure. Taking together the 6 data sets, a total of 1,908,092 particles were selected from 13,484 micrographs. Particles from each data set were separately subjected to 2 rounds of 2D classification in RELION2 to remove junk particles and contaminations resulting in a total of 505,681 particles for further processing (Supplementary Fig. 2b). Particles from the 6 data sets were merged into 2 independent data sets that were each subjected to 2 rounds of 3D *ab-initio* classification in cryoSPARC ^28^ with a class probability threshold of 0.9. After discarding the poor-quality models, the remaining particles were merged, resulting in a final data set of 162,332 particles. This final data set was used to generate a high-quality *ab-initio* model with cryoSPARC (Supplementary Fig. 2c). The *ab-initio* model was low-pass filtered to 30Å and was used as a reference for 3D auto-refinement in RELION2 producing a map of 7 Å global resolution. The refined particle coordinates were then used for local CTF estimation using GCTF v1.06 followed by a re-extraction with centering. This new particle stack was subjected to a 3D auto-refinement in RELION2 using the *ab-initio* model low-pass filtered at 30 Å resulting in a 6.8 Å map, further post-processed to a 6.6 Å global resolution (Supplementary Fig. 3). Local resolution calculated with Blocres ^55^ showed a range of resolution from 4Å in the DNA-binding/cleavage domain and 14Å in the ATPase domain indicating a high flexibility of the head. Moreover, 2D classifications showed large movements of the ATPase domain relative to the DNA-binding/cleavage domain (Supplementary Fig. 1d).

A combination of local approaches was used to identify different conformations and to improve local resolution of each domain. Since the ATPase head domain was too small for a 3D focused refinement, we first performed a focused 3D classification of the ATPase domain with a soft mask and no alignment in RELION2. One class of 36,610 particles with well-defined densities of the ATPase domain was selected. To facilitate an accurate alignment of the particles, a new *ab-initio* model of the class was calculated in cryoSPARC together with the DNA-binding/cleavage domain. Homogeneous refinement in cryoSPARC of the selected class yielded a reconstruction with overall resolution of 7.6Å in which density of the linkers appeared (Supplementary Fig. 3).

Secondly, a focused 3D auto-refinement was performed in RELION2 using a soft mask around the DNA-binding/cleavage domain. Post-processing of the map produced a 4.5Å resolution reconstruction of the DNA-binding/cleavage domain. Then, a focused 3D classification of the DNA-binding/cleavage domain starting with a fine angular sampling of 3.7°, local angular search range of 5° and tau fudge of 16 was performed. Two of the classes showed better angular accuracies and distinct conformations of the DNA-binding/cleavage domain, referred to as State 1 (57,976 particles) and State 2 (34,922 particles). These 2 classes were further refined in cryoSPARC by homogeneous refinement with a C2 symmetry and gave a reconstruction with global resolution of 4.2 Å and 5.0 Å, respectively. Reconstructions with relative positions of the ATPase domain regarding both State 1 and State 2 conformations of the DNA-binding/cleavage domain were obtained by a homogeneous refinement in cryoSPARC with a C1 symmetry. Auto-masking was disabled to avoid appearance of artifacts during refinement. The reconstruction of the overall complex with the DNA-binding/cleavage domain in State 1 conformation yielded a map of 5.6 Å overall resolution. For the DNA-binding/cleavage domain in State 2 conformation, the reconstruction yielded a map of the entire complex at 7.2Å overall resolution. Further heterogeneous refinements were performed to assess the flexibility of the ATPase domain with respect to the DNA-binding/cleavage domain (Supplementary Fig. 3).

All reported resolutions are based on the gold standard FSC-0.143 criterion ^56^ and FSC-curves were corrected for the convolution effects of a soft mask using high-resolution noise-substitution in RELION2 as well as in cryoSPARC ^57^. All reconstructions were sharpened by applying a negative B-factor that was estimated using automated procedures ^58^. Local resolution of maps reconstructed in cryoSPARC were calculated using Blocres (Supplementary Fig. 4). The cryo-EM maps of the DNA-binding/cleavage domain in State 1 and State 2 state as well as the entire complex in the 2 same conformations have been deposited in the EMDataBank under accession numbers EMD-11550, EMD-11551, EMD-11553, EMD-11554, respectively. The EM map with well-resolved N-gate/DNA-gate linkers have also been deposited in the EMDataBank under accession numbers EMD-11552.

### Model building and refinement of the DNA-binding/cleavage domain

The 2 reconstructions of the DNA-binding/cleavage domain in State 1 and State 2 at 4.2Å and 5.0Å, respectively, were used to refine a crystal structure of the hTop2α DNA-binding/cleavage domain in complex with DNA and etoposide ^29^. PDB 5GWK was stripped of all ions and water molecules, with all occupancies set to 1 and B-factors set to 50. First, the atomic model was rigid-body fitted in the filtered and sharpened maps with Chimera ^59^. A first round of real-space refinement in PHENIX ^60^ was performed using local real-space fitting and global gradient-driven minimization refinement. Then, nucleic acids were modified according to the DNA sequence used in our structure. The visible part of the CTD linker (1187-1215) was built as a poly-A coil. Several rounds of real-space refinement in PHENIX using restraints for secondary structure, rotamers, Ramachandran, and non-crystallographic symmetry were performed, always followed by manual inspection in COOT ^61^, until a converging model was obtained. A half-map cross-validation was performed to define 0.5 as the best refinement weight in PHENIX allowing atom clashes reduction and prevention of model overfitting (Supplementary Fig. 4). All refinement steps were done using the resolution limit of the reconstructions according to the gold standard FSC-0.143 criterion ^56^. Refinement parameters, model statistics and validation scores are summarized in Supplementary Table 1. The atomic models of the DNA-binding/cleavage domain in State 1 and State 2 conformations have been deposited in the Protein Data Bank under accession numbers 6ZY5, 6ZY6, respectively.

### Model building and refinement of the overall complex

For both conformations of the DNA-binding/cleavage domain, the 3D reconstructions of the overall complex were used for further atomic model refinement. The atomic models previously refined for each conformation of the DNA-binding/cleavage domain were rigid-body fitted in the overall maps using Chimera. Then, crystal structure of the ATPase domain in complex with ADPNP was rigid-body fitted in the filtered and unsharpened maps using Chimera. PDB 1ZXM ^8^ was stripped of all ions and water molecules, with all occupancies set to 1 and B-factors set to 50. A first round of real-space refinement in PHENIX was performed using rigid-body and global gradient-driven minimization refinement. Then, the linker between ATPase domain and the DNA-binding/cleavage domain was built in COOT as an alpha helix, following the density and according to the secondary structure prediction (Supplementary Fig. 6). Refinement followed the same procedure as for the masked DNA-binding/cleavage domain except that the local real-space fitting was replaced by a rigid-body refinement. Resolution limit for refinements was set according to the gold standard FSC-0.143 criterion. Refinement parameters, model statistics and validation scores are summarized in Supplementary Table 1. The atomic models of the full-length hTop2α in State 1 and State 2 conformations have been deposited in the Protein Data Bank under accession numbers 6ZY7, 6ZY8, respectively.

### Expression and purification of hTop2α mutants

The modified pVote0GW vector used for wild-type hTop2α overexpression was mutated by site directed mutagenesis using the QuikChange XL Site-Directed Mutagenesis kit (Agilent) in order to generate the plasmids harboring K418A, W414A-F417S, K425G-K426G-C427P mutations and Δ1193 or Δ1217 truncations (Supplementary Table 3). The overexpression procedure for the five mutants is identical to the wild type hTop2α described above in the Methods section. Purification of each mutant was performed in batch. Cells were harvested and resuspended in Lysis buffer (25 mM Hepes, 500 mM NaCl, 500 mM KCl, 1 mM MgCl_2_, 20 mM imidazole, 10% v/v glycerol, 2.5 mM beta-mercaptoethanol, 0.5 mM PMSF, 0.5 mM Pefabloc, protease inhibitor cocktail (Roche), pH 8.0) and lysed with 3 cycles of high pressure disruption using EmulsiFlex-C3 (Avestin) at 1500 bars. After centrifugation 1h at 40000g, the lysate of the 3 mutants (K418A, W414A-F417S, K425G-K426G-C427P) were incubated with 2 ml of TALON resin for 2h under agitation at 4°C. The proteins were washed 3 times with the lysis buffer containing increasing concentration of imidazole up to 50mM. Then proteins were eluted from the resin with 250mM imidazol pH 8.0 added to the Lysis buffer. The three aforementioned eluted proteins and the 2 additional mutants lysates (Δ1193 and Δ1217) were incubated with 2 ml of StrepTactin resin and incubated 2h under agitation at 4°C. The proteins were washed 3 times with 25 mM Hepes, 200 mM NaCl, 200 mM KCl, 1 mM MgCl_2_, 10% v/v glycerol, 2 mM DTT, pH 8.0 and eluted with the same buffer supplemented with 3 mM Desthiobiotin (Sigma).

### Relaxation assay

An increasing concentration of hTop2α was incubated at 37°C with 6 nmoles of supercoiled pUC19 plasmid in a reaction mixture containing 20 mM Tris-acetate pH7.9, 100 mM potassium acetate, 10 mM magnesium acetate, 1mM DTT, 1 mM ATP, 100 μg/ml BSA. After 30 minutes, reactions were stopped by addition of SDS 1%. Agarose gel electrophoresis was used to monitor the conversion of supercoiled pUC19 to the relaxed form. Samples were run on a 0.8% agarose, 1X Tris Borate EDTA buffer (TBE) gel, at 6V/cm for 180 min at room temperature. Agarose gels were stained with 0.5mg/ml ethidium bromide in 1X TBE for 15 min, followed by 5 min destaining in water. DNA topoisomers were revealed using a Synergy U:Genius 3 scanner.

### Cleavage assay

An increasing concentration of hTop2α was incubated at 37°C with 6 nmoles of supercoiled pUC19 plasmid in a reaction mixture containing 20 mM Tris-acetate pH7.9, 50 mM potassium acetate, 10 mM magnesium acetate, 1 mM DTT, 1 mM ATP, 100 μg/ml BSA and 275 μM of etoposide. After 30 minutes, reactions were stopped by addition of SDS 1% and 650 μM of proteinase K. After incubation at 37°C 40 minutes, a phenol-chloroform DNA extraction at pH 8 was performed. Agarose gel electrophoresis was used to monitor the conversion of supercoiled pUC19 to the relaxed form. Samples were run on a 0.8% agarose, 1X Tris Borate EDTA buffer (TBE) gel, at 6V/cm for 180 min at room temperature. Agarose gels were stained with 0.5 mg/ml ethidium bromide in 1X TBE for 15 min, followed by 5 min destaining in water. DNA topoisomers were revealed using a Synergy U:Genius 3 scanner.

### ATPase enzymatic assays

ATP hydrolysis was measured by following the oxidation of NADH mediated by pyruvate kinase (PK) and lactate dehydrogenase (LDH). The absorbance is monitored at 340 nm over 200 seconds at 37 °C with a Shimadzu 1700 spectrophotometer. Reactions were recorded in triplicates with 75 nM protein and 21 nM plasmid DNA (PUC19) in 500 μl of a buffer containing 50 mM Tris-HCl pH7.5, 150 mM potassium acetate, 8 mM magnesium acetate, 7 mM BME, 100 μg/mg of BSA, 4U/5U of PK/LDH mixture, 2 mM PEP, and 0.2 mM NADH and 4 mM ATP.

### Phylogenetic and structure prediction analysis of the TOP2

41 TOP2 genes, from unicellular eukaryotes to Human (Supplementary Table 4), were aligned with ClustalW ^62^ using the BLOSUM weight matrix. The subsequent alignment was injected into the ConSurf server ^63^ to analyze the conservation of the primary sequence. Secondary structure prediction of the hTop2α was performed using the PSIPRED server ^64^. Protein domain graphs (Figures 4 and 5) were generated using domainsGraph.py (archived at https://github.com/elifesciences-publications/domainsGraph) ^65^.

### hTop2α CTD production and purification

The sequence coding for the human Top2α CTD (1191-1531) was inserted into a modified pAX vector containing an N-terminal 10 His-tag and a C-terminal twin-strep tag. Overexpression was performed in *E. coli* BL21 (DE3) pRARE2 in LB medium containing 50 μg/mL kanamycin and 35 μg/mL chloramphenicol. Cells were induced with 1 mM IPTG after reaching an OD_600_ 0.95 and protein was expressed at 37 °C for 4 h. The CTD was purified similarly as described for the full length human Top2a supplemented by an additional size-exclusion chromatography step. After the ion exchange chromatography, fractions containing the intact CTD were pooled and loaded on a Superdex S200 16/60 size-exclusion chromatography column (GE Healthcare) using 25 mM Hepes, 200 mM NaCl, 200 mM KCl, 1 mM MgCl_2_, 10% v/v glycerol, 2 mM DTT, pH 8.0. After the purification process, 4 mg of the hTop2α CTD were obtained per liter of cell culture (Supplementary Fig. 9a).

### SAXS experiments

Small Angle X-ray Scattering (SAXS) data of the hTop2α CTD (1191-1531) at 10 or 20 mg/ml was collected on an in-house Rigaku BioSAXS-1000 (Rigaku) apparatus, equipped with a Rigaku MicroMaxTM-007HF generator and a Pilatus 100k detector. The sample was maintained at 10°C and exposed to the X-ray beam for 2h. A total of 8 images were recorded every 15 min. All the data processing steps were performed using the program package PRIMUS ^66^.

