## Supplementary information for "Structural basis for the allosteric regulation of Human Topoisomerase 2α"

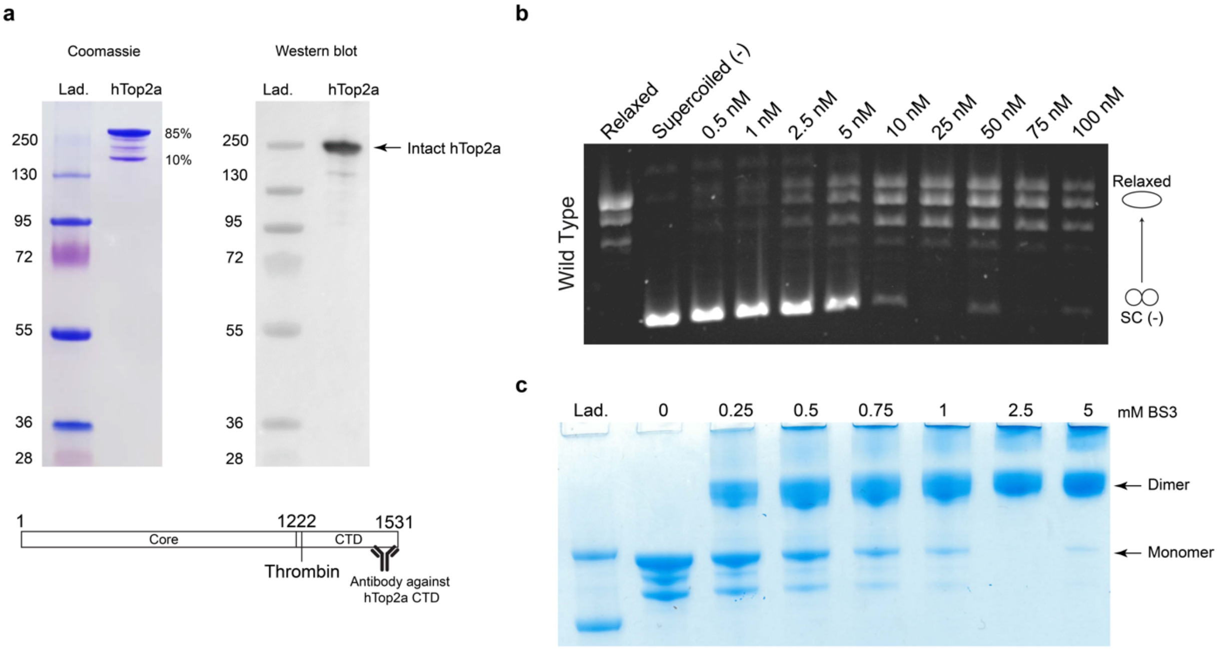


**Supplementary Fig. 1. Purification and stabilization of the hTop2α. a.** SDS-PAGE and Western blot analysis of the purified hTop2α. Coomassie staining is shown on the left. Western blot analysis using a monoclonal antibody directed against the hTop2α CTD is shown on the right. During the protein preparation, the CTD is cleaved off in 10 to 15% of the sample. **b.** Relaxation activity by wildtype hTop2α. Protein concentrations are listed in nM of holoenzyme. **c.** Titration of BS3 for stabilization of the DNA-bound hTop2α homodimer. Formation of the full-length complex can still be obtained from the predominant species in presence of DNA and after crosslinking with BS3.

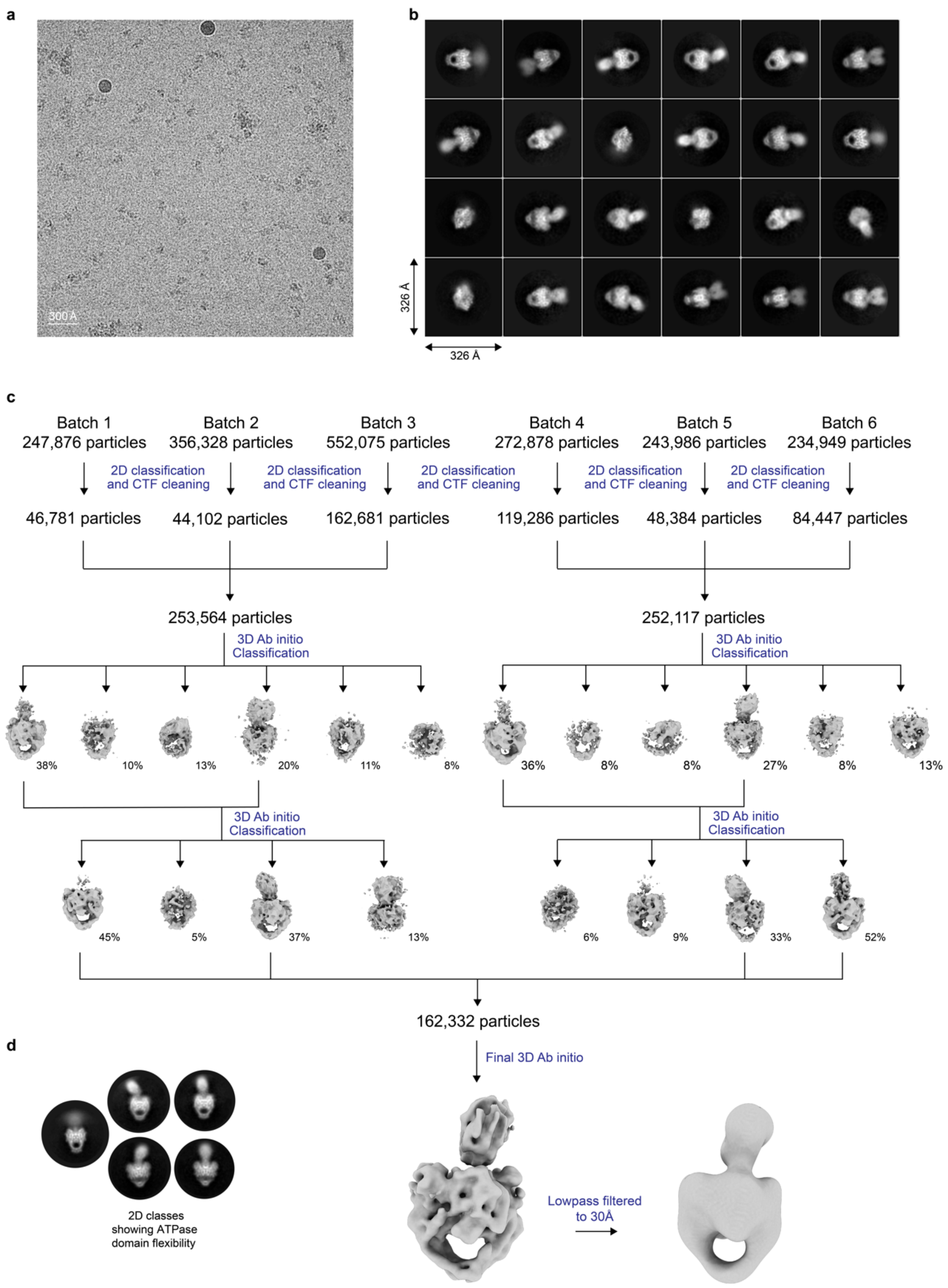


**Supplementary Fig. 2. Cryo-EM data acquisition and *ab-initio* model generation. a.** A typical Cryo-EM micrograph collected on a FEI Titan Krios microscope operated at 300 kV and detected with a Gatan K2 Summit camera. **b**. Reference-free 2D classification. **c**. Flow chart of data processing from 2D classification to *ab-initio* model generation. The number of particles is indicated at each step. **d.** Subset selection of 2D classes showing the flexibility of the ATPase domain with respect to the DNA-binding/cleavage domain.

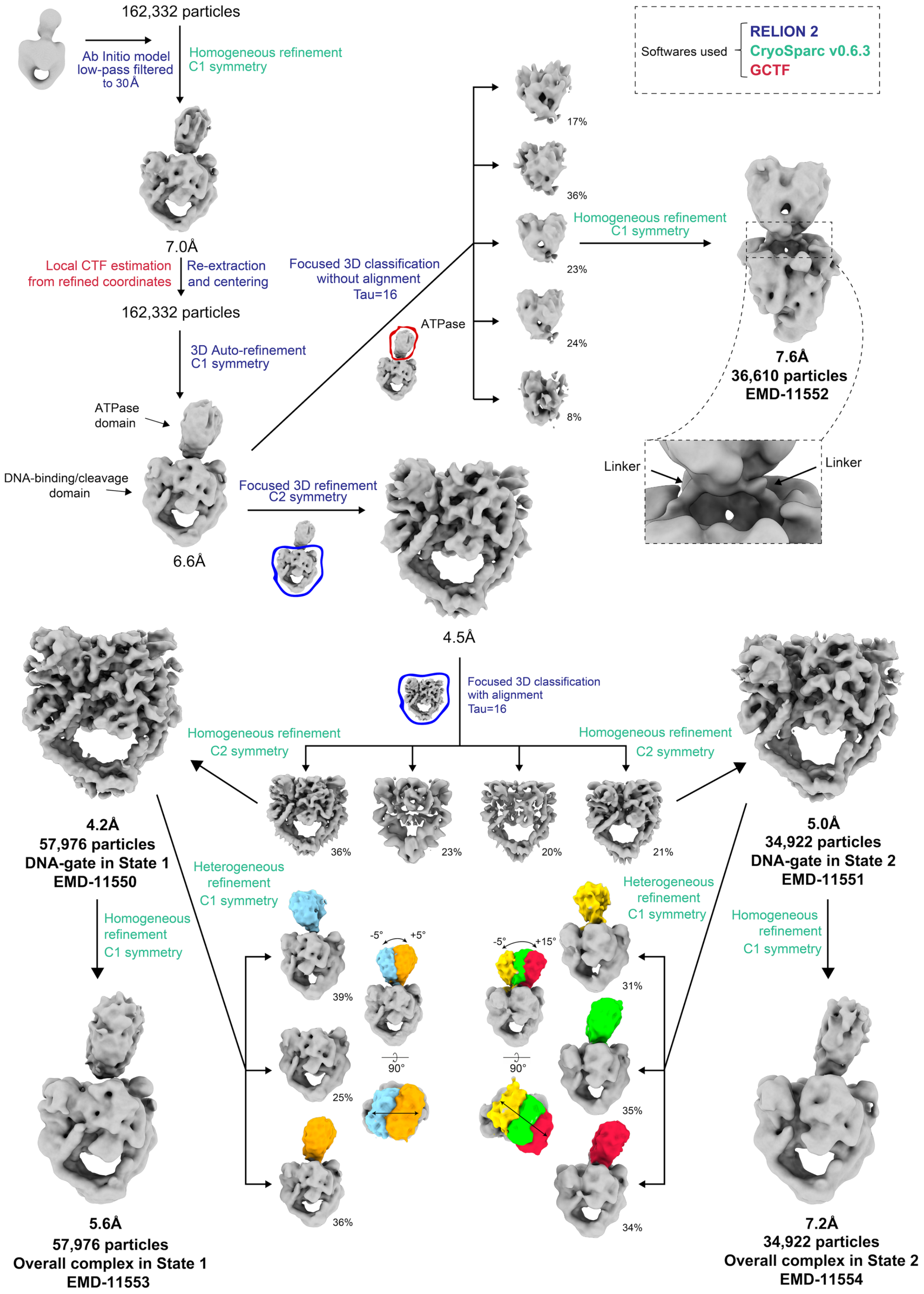


**Supplementary Fig. 3. Flow chart of cryo-EM data processing.** The *ab-initio* model was refined in cryoSPARC yielding a map of 6.6Å overall resolution using 162,332 particles. Focused 3D classification with and without alignment followed by Homogeneous refinement allowed us to solve 4 new structures: the DNA-binding/cleavage core in a closed state at 4.2Å resolution and in a pre-opening state at 5.0Å resolution, the overall complex in a closed state at 5.6Å resolution and the overall complex in pre-opening state at 7.2Å resolution. The number of particles used for the final refinement is indicated.

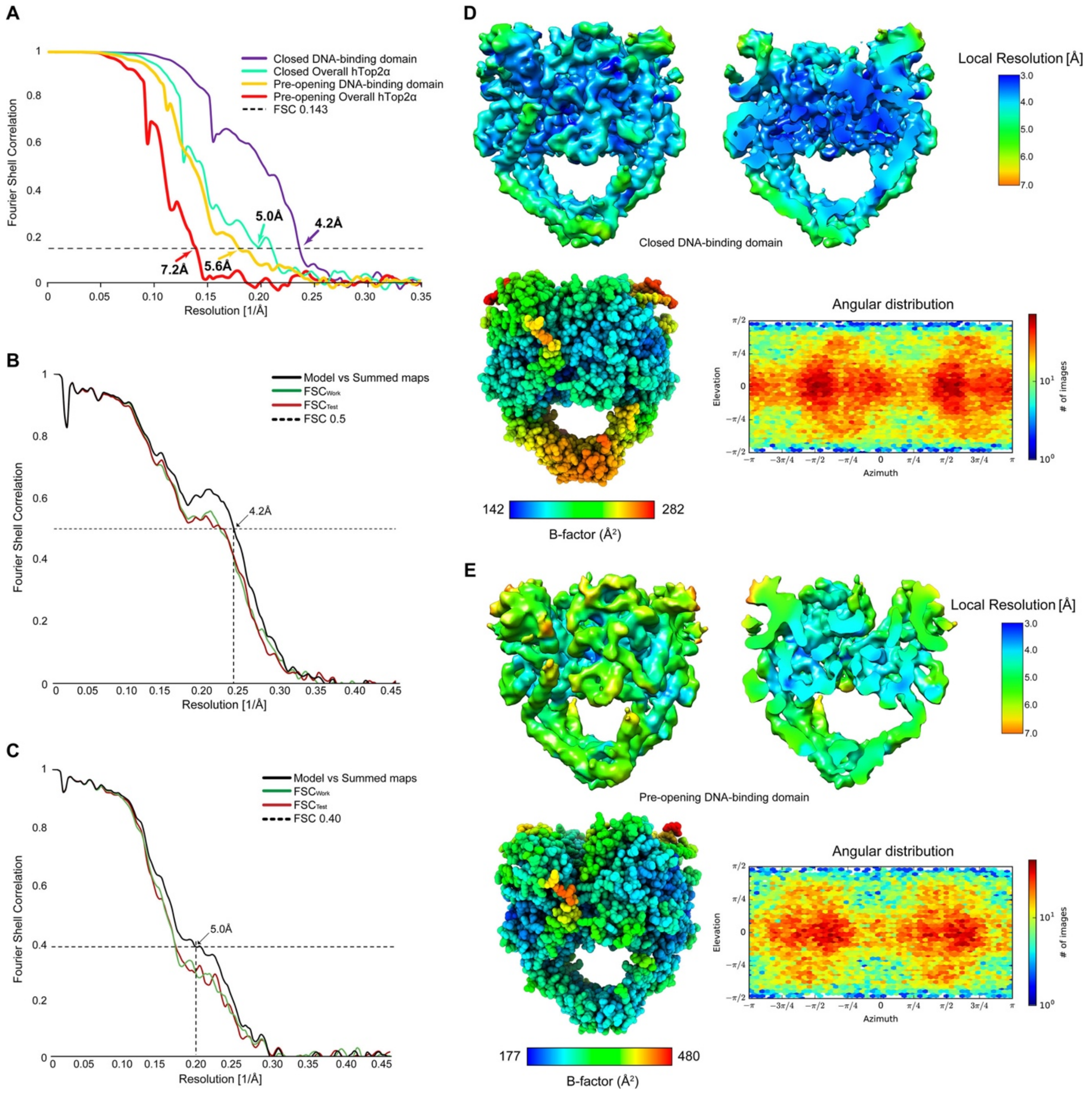


**Supplementary Fig. 4. Cryo-EM statistics of the overall DNA-bound hTop2α and DNA-binding/cleavage domain in closed and pre-opening states. a.** FSC plots and resolution estimation using the gold-standard 0.143 criterion generated from RELION2. Cross-validation FSC curves for the closed (**b**) and pre-opening (**c**) DNA-binding/cleavage domain models versus unfiltered half maps (the one used in the refinement, $FSC_{work}$, and the other half, $FSC_{free}$) and the unfiltered summed maps. Final refined map of the closed (**d**) and pre-opening (**e**) DNA-binding/cleavage domain. The maps (upper left) and sliced maps (upper right) are coloured according to local resolution calculated with Blocres. The atomic models refined in the corresponding cryo-EM maps (lower left) are colored according to the B-factors. The angular distribution plots (lower right) were generated from cryoSPARC.

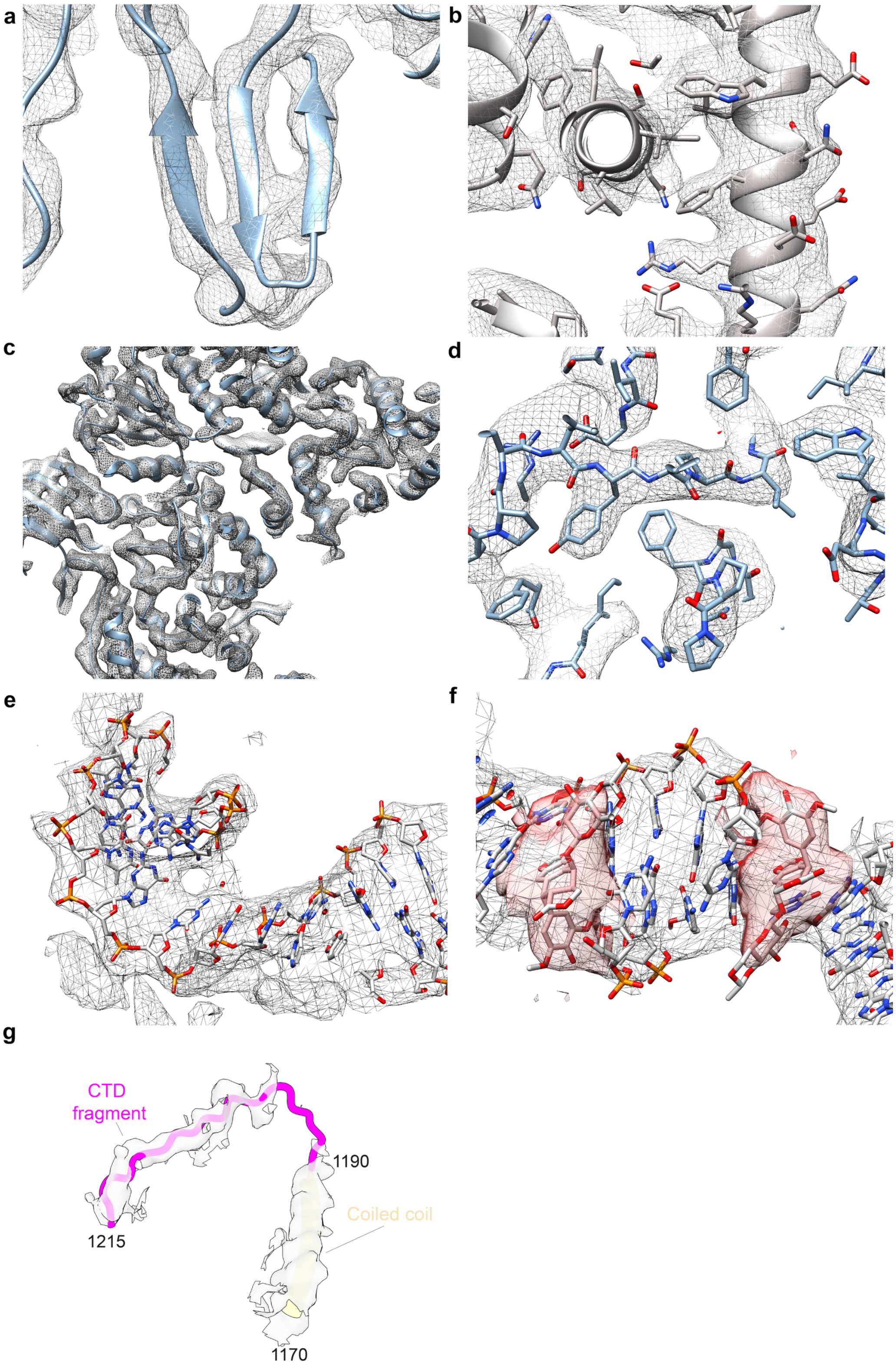


**Supplementary Fig. 5. Near-atomic resolution features of the DNA-binding/cleavage domain in closed conformation.** a. Beta strands individualization. **b.** Alpha helices with well-defined EM density of the side chains. **c.** Overview of an atomic model region fitted in the EM density. **d.** Aromatic residues with well-defined EM density. **e.** G-segment fitted in the EM density. **f.** EM density of etoposide in red. **g.** EM density of the CTD linker (magenta line, residues 1191-1215).

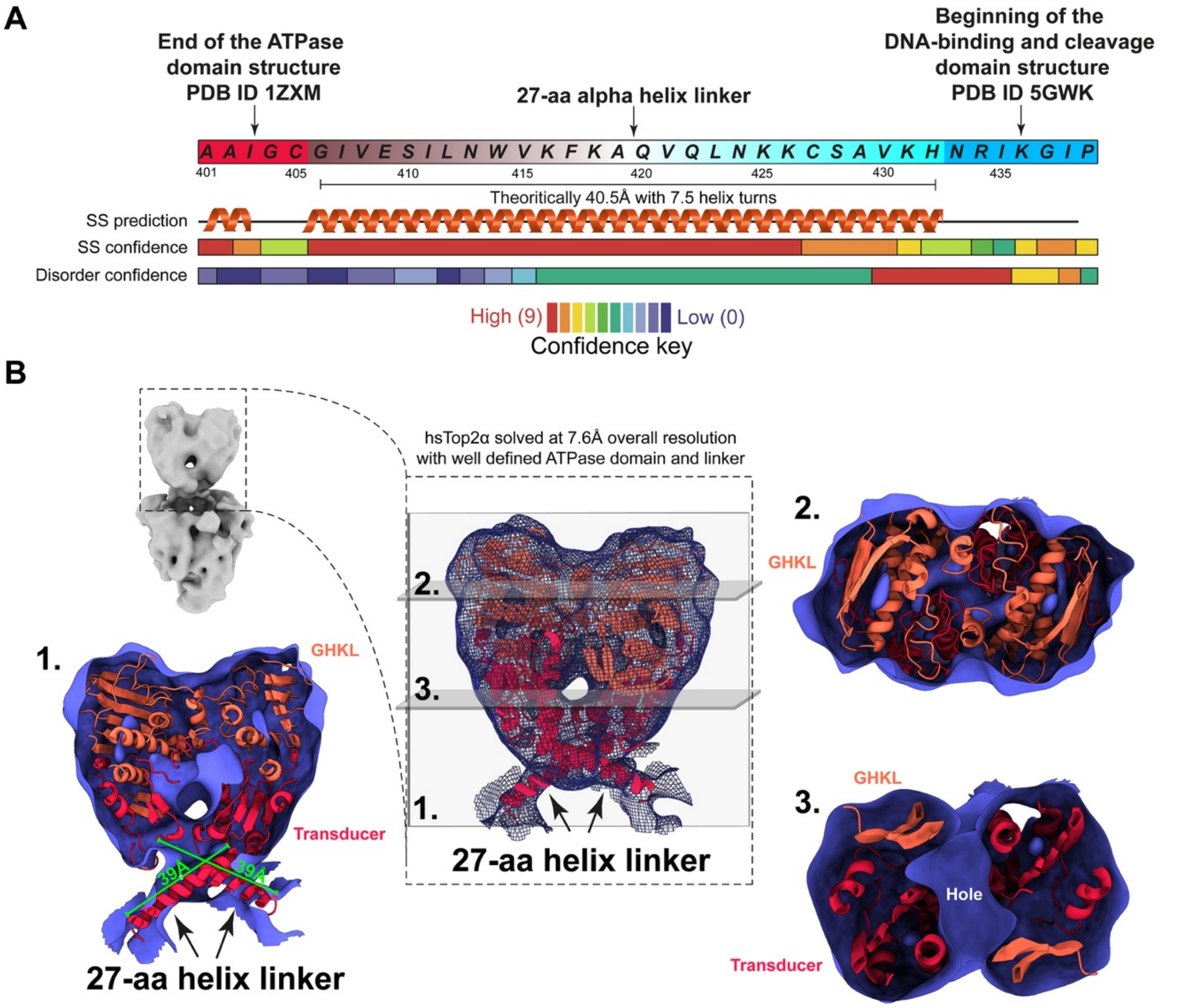


**Supplementary Fig. 6. Building of the 27-aa linkers between the N-gate and the DNA gate. a.** The 27-aa linkers are predicted to fold as an alpha helix based on secondary structure prediction performed with Phyre2 [1]. The theoretical length of a 27-aa alpha helix is 40.5Å with 7.5 turns, considering that an alpha helix has 3.6 residues per turn and a pitch length of 5.4Å. **b.** After fitting of the functional domains in the EM density, the distance between the last residue of the ATPase domain, C405, (PDB ID 1ZXM) and the first residue of the DNA-binding/cleavage domain, N433, is 39Å. To accommodate such distance with the missing 27-aa, the linkers were built as alpha helices accordingly to the secondary structure prediction and the EM density map. The 3 views of the ATPase domain through different slices of EM density (7.6 Å resolution) show an overall good fit of the atomic model in the map.

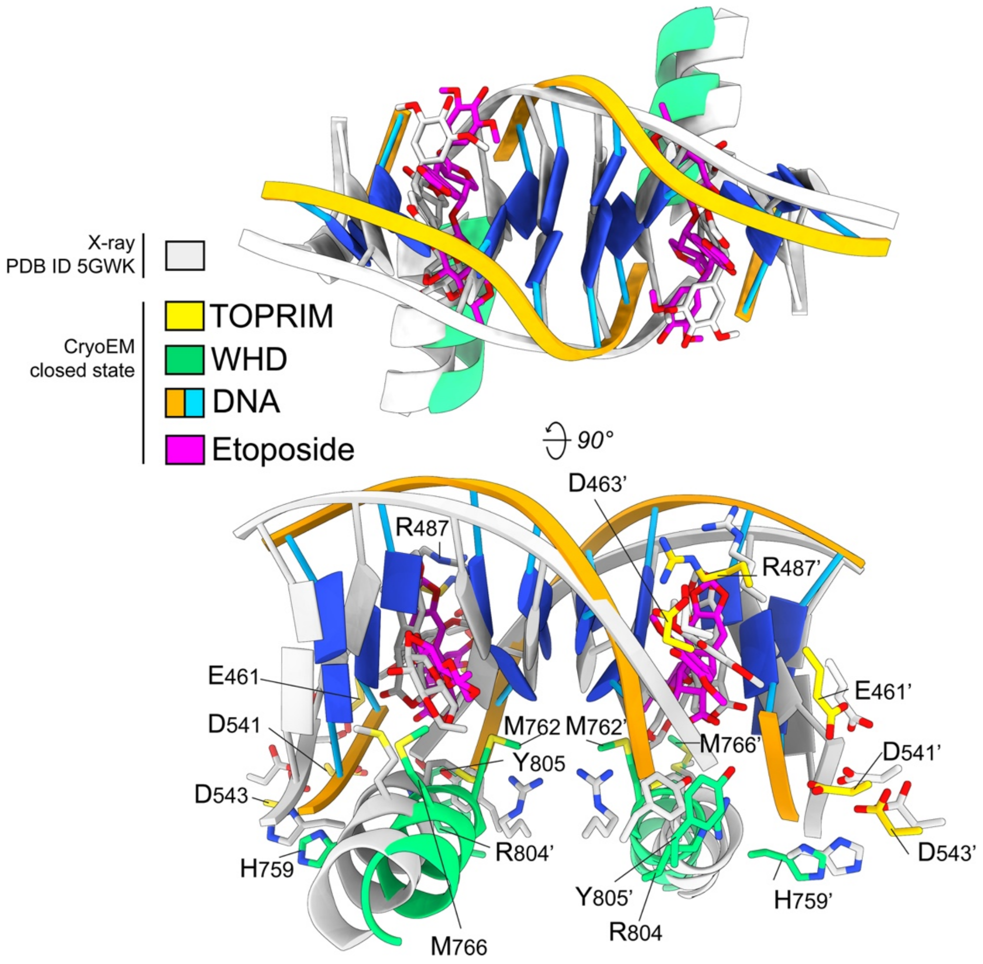


**Supplementary Fig. 7. Comparison of etoposide binding sites of cryo-EM and X-ray hTop2α structures.** Slight rearrangement of the protein structure and DNA bases in the EM structure, compared to the X-ray structure, yielded to a minor shift of the etoposide in the binding site during refinement to avoid steric clashes. Structures are colored as indicated in the legend.

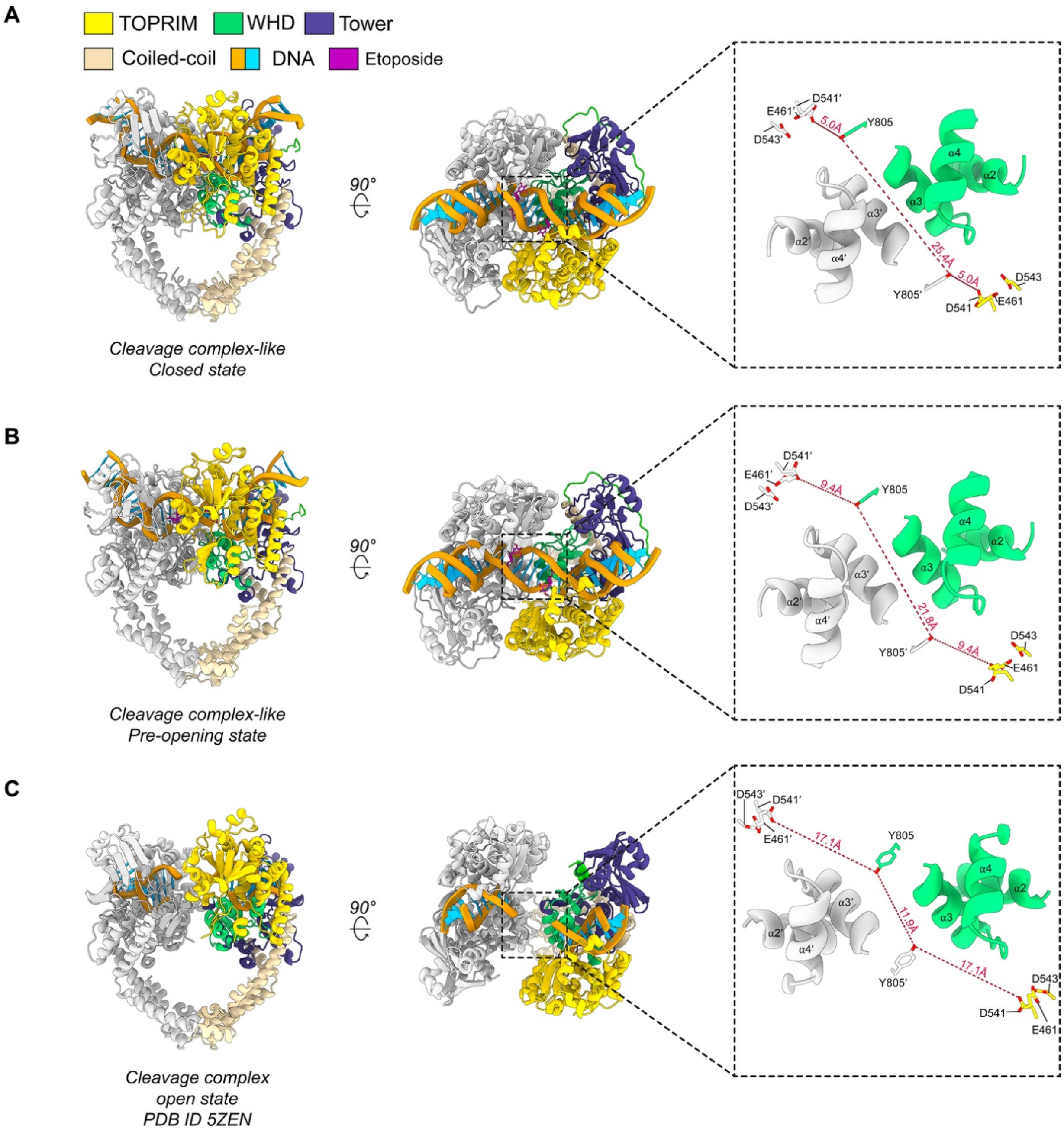


**Supplementary Fig. 8. Quaternary and tertiary changes associated with G-segment opening after cleavage.** Orthogonal views of the hTop2α DNA-binding/cleavage core in different states. The inset show<s selected structural elements (alpha helices α2, α3 and α4) lining the bottom side of the G-segment groove and key catalytic amino acids in the closed cleavage complex-like (CC-like) (**a**), pre-opening CC-like (**b**) and open CC (**c**) [2]. For each conformation, the catalytic tyrosine residues, the cation-binding residues and the distances between the two catalytic residues and the closer aspartic acid residue of the DxD dyad are shown to illustrate structural changes in the DNA-gate during closed-to-open transitions. See legend on the top for the color code. Residues from different homodimer are colored differently, with labels belonging to the second homodimer marked by a prime.

**Supplemental analysis related to Supplementary Fig. 8**

The main conformational changes are observed in the DNA-gate, while the ATPase domain is more prone to rotations and translational movements to accompany the DNA-gate motions (Figure 3). During this conformational transition, the two alpha helices α3 and α3' of each homodimer slide against each other by half helix turn yielding the catalytic tyrosine residues rapprochement by 3.6 Å. As the TOPRIM domain performs a swinging outward movement, the DxD di-acidic metal ion-binding motif recede by 4.4 Å from the catalytic tyrosine residues, decoupling the key catalytic residues which disfavor the religation of cleaved DNA ends (Supplementary Fig. 8a-b). Concomitantly to the motion of the DNA-Gate, the dimerized ATPase domain is rotating by 3° counterclockwise (opposite to the intertwining) and is coming closer to the DNA-gate in order to push the T-segment in the newly formed groove between the TOPRIM and tower domains (Figure 3). These two states of the DNA-gate precede the open conformation [2]. In particular, the two alpha helices α3 and α3' of each homodimer have slid against each other by one helix turn, bringing closer the catalytic tyrosine residues. The DxD dyads are now far from the tyrosine residues and follow the sliding and swiveling motion of the DNA-gate (Supplementary Fig. 8c).

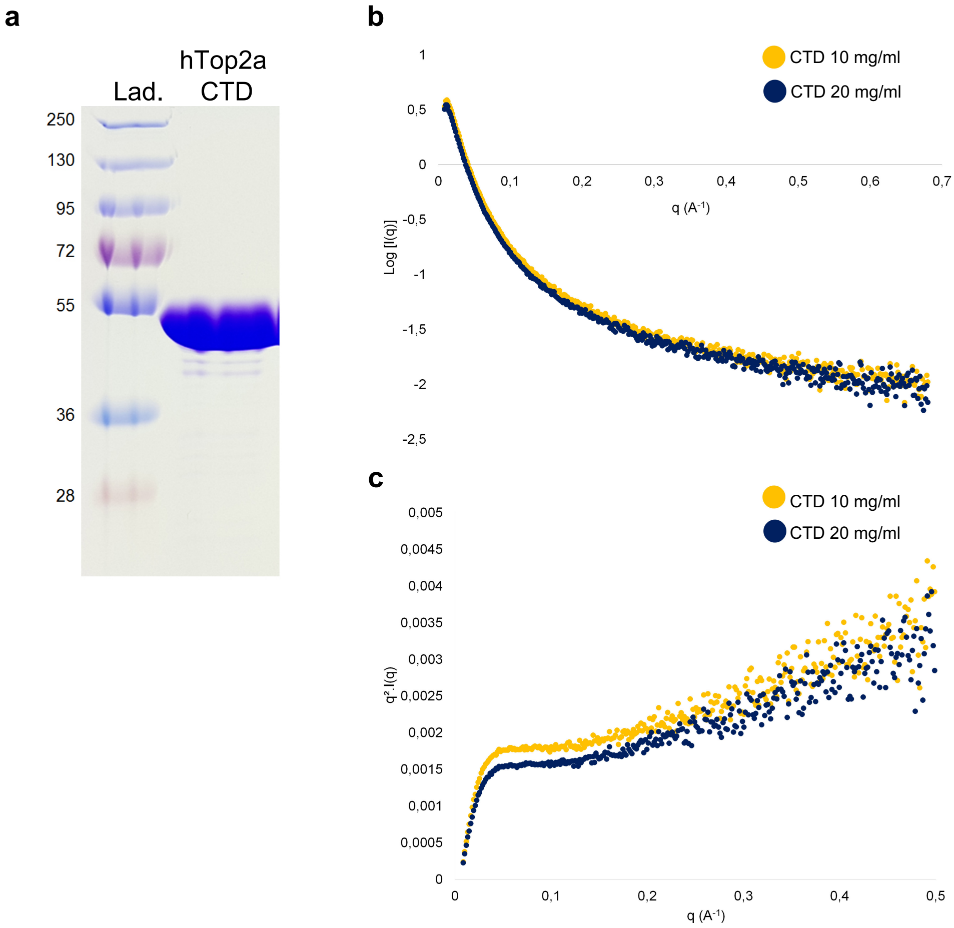


**Supplementary Fig. 9. SAXS analysis of the hTop2α CTD. a.** SDS-PAGE analysis of the purified hTop2α CTD (1191-1531) **b.** Experimental SAXS curves of the CTD at 10 mg/ml (yellow) and 20 mg/ml (blue). **c.** Kratky plot demonstrating the absence of fold of the CTD.

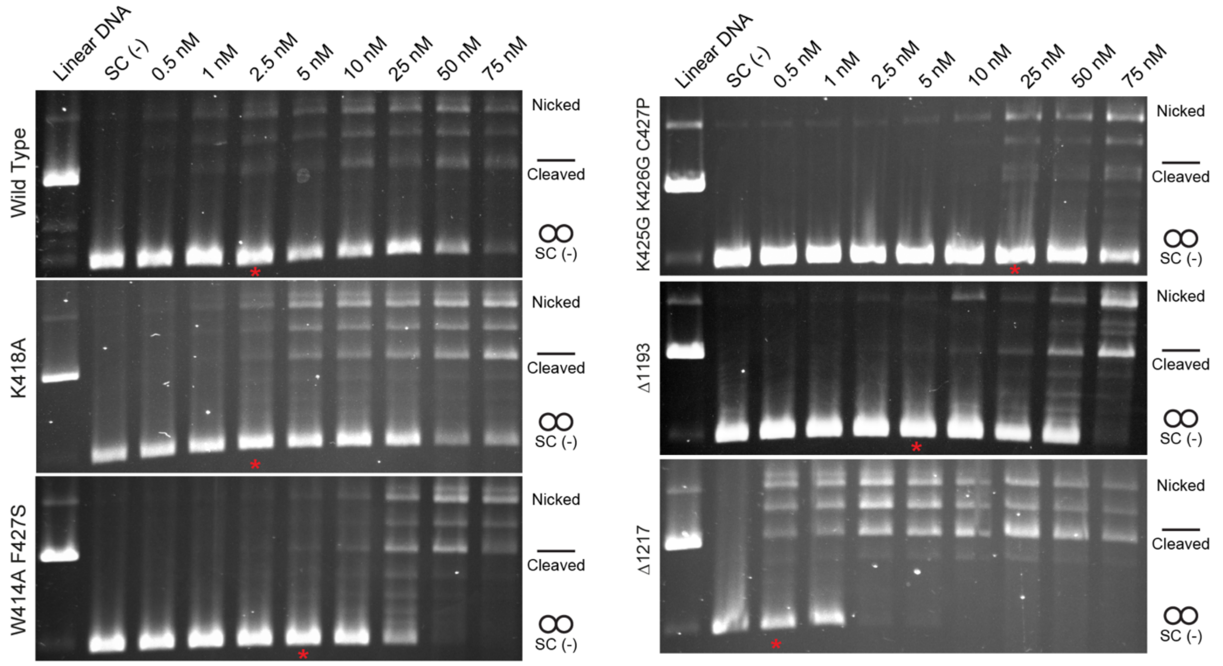


**Supplementary Fig. 10. DNA Cleavage assays.** Comparison of cleavage activity in presence of 250 µM etoposide of wild-type, K418A, W414A-F417S and K425G-K426G-C427P, Δ1193 and Δ1217 hTop2α. Protein concentrations are listed in nM of holoenzyme. The asterisks indicate the lowest concentration at which a cleavage band is observed.

| **Data Collection** | | | | | | |
|---|---|---|---|---|---|---|
| Microscope | Titan Krios | | | | | |
| Voltage (keV) | 300 | | | | | |
| Magnification | 105,000 | | | | | |
| Electron dose ($e^{-}$ Å$^{-2}$) | 50 | | | | | |
| Detector | Gatan K2 Summit (super-resolution mode) | | | | | |
| Pixel Size (Å) | 1.1 (0.55) | | | | | |
| Defocus range (µm) | -1 to -3 | | | | | |
| **Individual data sets** | | | | | | |
| Batches | **1** | **2** | **3** | **4** | **5** | **6** |
| Micrographs (no.) | 2299 | 2230 | 3128 | 2348 | 1553 | 1926 |
| Extracted particles (no.) | 247,876 | 356,328 | 552,075 | 272,878 | 243,986 | 234,949 |
| Particles after cleaning (2 rounds of 2D classification, MaxRes<5Å) | 46,781 | 44,102 | 162,681 | 119,286 | 48,384 | 84,447 |

| **Merged data sets for refinement** | |
|---|---|
| Total merged particles | 505,681 |
| Particles after cleaning (2 rounds of 3D ab initio classification) | 162,332 |

| **Reconstructions** | | | | |
|---|---|---|---|---|
| | DNA-binding/cleavage domain in state 1 | DNA-binding/cleavage domain in state 2 | Overall complex in state 1 | Overall complex in state 2 |
| | EMD-11550 | EMD-11551 | EMD-11553 | EMD-11554 |
| | PDB 6ZY5 | PDB 6ZY6 | PDB 6ZY7 | PDB 6ZY8 |
| Software | cryoSPARC v0.6.3 | | | |
| Final particles (no.) | 57976 | 34922 | 57976 | 34922 |
| Box size (pixels) | 296x296x296 | | | |
| Symmetry imposed | C2 | C2 | C1 | C1 |
| Map resolution FSC 0.143 (global) (Å) | 4.22 | 5.04 | 5.56 | 7.18 |
| Applied B-factor for sharpening (Å$^2$) | -165 | -295 | -107 | -161 |

| **Model refinement** | | | | |
|---|---|---|---|---|
| Software | Phenix 1.12 | | | |
| Resolution cut-off (Å) | 4.2 | 5.0 | 5.6 | 7.2 |
| Non-hydrogen atoms | 13510 | 13510 | 19985 | 19980 |
| Protein residues | 1520 | 1520 | 2319 | 2318 |
| DNA bases (atoms) | 1218 | 1218 | 1218 | 1218 |
| Ligands (atoms) | 84 | 84 | 146 | 146 |
| Average B-factor | 190 | 264 | 635 | 762 |
| R.m.s. deviations | | | | |
| Bond lengths (Å) | 0.004 | 0.004 | 0.004 | 0.004 |
| Bond angles (°) | 0.832 | 0.834 | 0.890 | 0.901 |
| **Validation** | | | | |
| Real space correlation coefficient | 0.83 | 0.84 | 0.79 | 0.73 |
| MolProbity score | 1.61 | 1.54 | 1.37 | 1.54 |
| Clashscore (all atoms) | 6.47 | 6.77 | 6.70 | 7.41 |
| Poor rotamers (%) | 0.15 | 0 | 0 | 0 |
| Ramachandran plot | | | | |
| Favored (%) | 96.23 | 97.09 | 98.05 | 97.27 |
| Allowed (%) | 3.77 | 2.91 | 1.95 | 2.73 |
| Outliers (%) | 0 | 0 | 0 | 0 |

**Supplementary Table 1. Data collection, processing and refinement statistics.**

| | |
|---|---|
| 13bp | GAGGATGACGATG |
| 17bp | CGCGCATCGTCATCCTC |

**Supplementary Table 2. Asymmetric oligonucleotides sequences.**

| Primer name | Sequence |
|---|---|
| hTop2a-K418A-fw | AAGCATACTAAACTGGGTGAAGTTTGCGGCCCAAGTCCAG |
| hTop2a-K418A-rev | CTGGACTTGGGCCGCAAACTTCACCCAGTTTAGTATGCTT |
| htop2A-W414A-F417S-fw | GTGGTATTGTAGAAAGCATACTAAACGCGGTGAAGAGTAAGGCCCAAGTCCAG |
| htop2A-W414A-F417S-rev | CTGGACTTGGGCCTTACTCTTCACCGCGTTTAGTATGCTTTCTACAATACCAC |
| hTop2a-KKC425GGP-fw | GGTGAAGTTTAAGGCCCAAGTCCAGTTAAACGGGGGGCCTTCAGCTGTAAAACATAATAGAATCAAGGGA |
| hTop2a-KKC425GGP-rev | TCCCTTGATTCTATTATGTTTTACAGCTGAAGGCCCCCCGTTTAACTGGACTTGGGCCTTAAACTTCACC |
| hTop2a-Delta1193-fw | GATGAACAAGTCGGACTTGAAGTTCTGTTCCAGGGG |
| hTop2a-Delta1193-rev | CCCCTGGAACAGAACTTCAAGTCCGACTTGTTCATC |
| hTop2a-Delta1217-fw | GCCTTCTCCGCGTGGTCTGGAAGTTCTGTTCC |
| hTop2a-Delta1217-rev | GGAACAGAACTTCCAGACCACGCGGAGAAGGC |

**Supplementary Table 3. Primer sequences used for the plasmids mutagenesis.**

| Protein name | Organism | Uniprot ID |
|---|---|---|
| TOP2 | *Aedes albopictus* | A0A023EXD7 |
| TOP2A | *Ailuropoda melanoleuca* | G1LN78 |
| TOP2A | *Anas platyrhynchos* | U3IFZ2 |
| TOP2 | *Arabidopsis thaliana* | P30182 |
| TOP2 | *Ascaris suum* | F1KQV5 |
| TOP2 | *Biomphalaria glabrata* | A0A2C9K2L3 |
| TOP2 | *Caenorhabditis elegans* | Q23670 |
| TOP2 | *Callorhinchus milii* | A0A4W3J3D7 |
| TOP2A | *Cavia porcellus* | H0V8L7 |
| TOP2 | *Ciona intestinalis* | F7AKG2 |
| TOP2 | *Crassostrea gigas* | K1Q404 |
| TOP2A | *Danio rerio* | Q6DRC7 |
| TOP2 | *Daphnia magna* | A0A165ADD4 |
| TOP2 | *Dictyostelium discoideum* | Q55BP5 |
| TOP2 | *Drosophila melanogaster* | P15348 |
| TOP2 | *Echinococcus granulosus* | W6UGB7 |
| TOP2A | *Gallus gallus* | O42130 |
| TOP2B | *Gallus gallus* | O42131 |
| TOP2A | *Homo sapiens* | P11388 |
| TOP2B | *Homo sapiens* | Q02880 |
| TOP2A | *Latimeria chalumnae* | H3ALH9 |
| TOP2 | *Loa loa* | A0A1I7VLJ5 |
| TOP2A | *Loxodonta africana* | G3UIA0 |
| TOP2A | *Monodelphis domestica* | F7ABY1 |
| TOP2A | *Mus musculus* | Q01320 |
| TOP2B | *Mus musculus* | Q64511 |
| TOP2 | *Octopus bimaculoides* | A0A0L8FFS9 |
| TOP2A | *Ophiophagus hannah* | V8P300 |
| TOP2A | *Pelodiscus sinensis* | K7F361 |
| TOP2 | *Saccharomyces cerevisiae* | P06786 |
| TOP2A | *Sarcophilus harrisii* | G3WFK3 |
| TOP2 | *Sarcoptes scabiei* | A0A132ALV4 |
| TOP2 | *Strongylocentrotus purpuratus* | W4XMF6 |
| TOP2 | *Stylophora pistillata* | A0A2B4S1C7 |
| TOP2 | *Tetrahymena thermophila* | Q6PUA4 |
| TOP2 | *Trypanosoma cruzi* | P30190 |
| TOP2A | *Tursiops truncatus* | A0A2U4BNZ1 |
| TOP2B | *Tursiops truncatus* | A0A2U4CFF8 |
| TOP2A | *Xenopus laevis* | Q6INT0 |

**Supplementary Table 4. Uniprot sequence ID of the homologs used for the multiple sequence alignments.**
